# CD147 antibody specifically and effectively inhibits infection and cytokine storm of SARS-CoV-2 variants

**DOI:** 10.1101/2021.05.14.444111

**Authors:** Jiejie Geng, Liang Chen, Yufeng Yuan, Ruo Chen, Ke Wang, Yongqiang Deng, Peng Du, Jiangning Liu, Guizhen Wu, Youchun Wang, Ke Xu, Xiuxuan Sun, Ting Guo, Xu Yang, Jiao Wu, Jianli Jiang, Ling Li, Jun Zhang, Kui Zhang, Hua Zhu, Zhaohui Zheng, Xianghui Fu, Fengfan Yang, Xiaochun Chen, Hao Tang, Zheng Zhang, Ding Wei, Yang Zhang, Ying Shi, Yumeng Zhu, Zhuo Pei, Fei Huo, Shirui Chen, Qingyi Wang, Wen Xie, Yirong Li, Mingyan Shi, Huijie Bian, Ping Zhu, Zhi-Nan Chen

## Abstract

SARS-CoV-2 and its variants are raging worldwide. Unfortunately, the global vaccination is not efficient enough to attain a vaccine-based herd-immunity and yet no special and effective drug is developed to contain the spread of the disease. Previously we have identified CD147 as a novel receptor for SARS-CoV-2 infection. Here, we demonstrated that CD147 antibody effectively inhibits infection and cytokine storm caused by SARS-CoV-2 variants. In CD147^KO^ VeroE6 cells, infections of SARS-CoV-2, its variants (B.1.1.7, B.1.351) and pseudovirus mutants (B.1.1.7, B.1.351, B.1.525, B.1.526 (S477N), B.1.526 (E484K), P.1, P.2, B.1.617.1, B.1.617.2) were decreased. Meanwhile, CD147 antibody effectively blocked the entry of variants and pseudomutants in VeroE6 cells, and inhibited the expression of cytokines. A model of SARS-CoV-2-infected hCD147 transgenic mice was constructed, which recapitulated the features of exudative diffuse alveolar damage and dynamic immune responses of COVID-19. CD147 antibody could effectively clear the virus and alveolar exudation, resolving the pneumonia. We found the elevated level of cyclophilin A (CyPA) in plasma of severe/critical cases, and identified CyPA as the most important proinflammatory intermediate causing cytokine storm. Mechanistically, spike protein of SARS-CoV-2 bound to CD147 and initiated the JAK-STAT pathway, which induced expression of CyPA. CyPA reciprocally bound to CD147, triggered MAPK pathway and consequently mediated the expression of cytokine and chemokine. In conclusion, CD147 is a critical target for SARS-CoV-2 variants and CD147 antibody is a promising drug to control the new wave of COVID-19 epidemic.

## Introduction

The rapid and continual mutation of SARS-CoV-2 is giving rise to a new wave of epidemic, with more than 150 million people infected globally, resulting in over 3 million deaths. Since the outbreak of the pandemic in 2020, virus variants have emerged globally, such as B.1.1.7 in the United Kingdom, B.1.351 in South Africa, B.1.526 in New York, P.1 in Brazil, and especially B.1.617 and B.1.618 discovered in India, which were reported to have double mutants and triple mutants respectively. These mutant strains have extensive mutations in spikes, including N501Y, E484K, E484Q, D614G, which accelerate the spread of the virus or facilitate to escape immune response (*1–3*). At present, global vaccination is the hope of eliminating the epidemic, but researchers have found that the existing vaccine is not effective against the mutant strains (*4*). Therefore, there is a need to develop safe and effective broad-spectrum approach that can control the rapid spread of SARS-CoV-2.

COVID-19 pneumonia patients, if not treated in a timely manner, deteriorate rapidly within 7 to 14 days, leading to acute respiratory distress syndrome, which represents the major cause of morbidity in COVID-19 patients (*5, 6*). The main pathological features of COVID-19 pneumonia comprise exudative diffuse alveolar damage with massive capillary congestion and hemorrhage (*7–9*). The pathology of these lung tissue damages, which has been described in up to 20% of COVID-19 patients, are thought to be related to cytokine storm syndrome (CSS). CSS induces exudative diffuse alveolar damage, fibrin exudation in the alveolar space, and infiltration of inflammatory cells such as macrophages and neutrophils, accompanied by diffuse hemorrhage and massive release of cytokines and chemokines. Immune analysis in COVID-19 patients highlighted profound impaired type I IFN responses characterized by a low level of IFN activity and downregulation of IFN stimulated genes, and an increased number of Th17 cells and their related inflammatory responses, leading to hyper-inflammation and acute lung injury (*10–12*). However, the mechanistical cause of CSS in COVID-19 pneumonia patients remains unclear due to a lack of suitable preclinical mouse model that recapitulate the pathology of COVID-19 pneumonia following SARS-CoV-2 infection.

To study the inflammatory responses in COVID-19, many laboratories have constructed mice expressed by human angiotensin-converting enzyme 2 (hACE2), which was introduced into the mice in different ways, such as transient introduction of hACE2 via adenoviral vectors (*13*) and expression of hACE2 driven with the mouse ACE2 (*14*), HFH4 (*15*) and K18 (*16*) promoters. These mouse models all supported SARS-CoV-2 infection and presented either mild or severe pneumonia features, including infiltration of inflammatory cells in the lung, high levels of proinflammatory cytokines and chemokines in lung homogenates, and severe interstitial and consolidative pneumonia (*17*). However, hACE2 mouse is not an ideal model to study the pathology of COVID-19 pneumonia, because SARS-CoV-2-infected hACE2 mice mostly exhibited interstitial pneumonia, and exudation was rarely observed, which cannot well simulate the pathological and immune characteristics of patients with COVID-19 pneumonia.

Our previous study discovered that CD147 is a novel receptor for SARS-CoV-2 infection, and human CD147 transgenic (hCD147) mice can be successfully infected with the virus (*18*). In this article, CD147^KO^ VeroE6 cells, SARS-CoV-2-infected hCD147 transgenic mice, and CD147 antibody were used to evaluate the variants infection and cytokine expression, and analyze the pathological phenotypes of pneumonia and immune characteristics. We found that the CD147 molecule was still a receptor for variants’ entry into host cells and could mediate cytokine storm. Thus, CD147 is confirmed to be a target for broad-spectrum of SARS-CoV-2 variants.

## Results

### CD147 antibody effectively inhibits the infection of SARS-CoV-2 variants

To determine whether CD147 is an infection target of SARS-CoV-2 variants, we used CRISPR-Cas9 to knock out CD147 on VeroE6 cells, and then carried out virus infection experiment using B.1.1.7 and B.1.351 variants. The results showed that the infection of B.1.1.7 and B.1.351 variants in VeroE6 cells was not significantly changed compared with that of SARS-CoV-2 (Fig. 1A). The knockout of CD147 decreased the infections of not only SARS-CoV-2 but also its two variants, B.1.1.7 and B.1.351 significantly (Fig. 1B). More importantly, CD147 antibody (Meplazumab) could effectively inhibit the infection of SARS-CoV-2, B.1.1.7 and B.1.351, at inhibition rates of 68.7%, 75.7% and 52.1% respectively for 60 μg/mL antibody (Fig. 1C). We further infected 9 strains of pseudoviruses mutants, including B.1.1.7, B.1.351, B.1.525, B.1.526(S477N), B.1.526(E484K), P.1, P.2, B.1.617.1, and B.1.617.2. The results showed that the infection of pseudoviruses was markedly suppressed by CD147 knockout (Fig. 1D), and CD147 antibody could effectively inhibit the infection of these pseudoviruses (Fig. 1E). B.1.1.7, B.1.351, B.1.525, B.1.526(S477N), B.1.526(E484K), P.1 and P.2 variants were inhibited by Meplazumab, and B.1.617.1 and B.1.617.2 variants were inhibited by a combination of Mehozumab (*China patent, Number: 201910796766.8*) and anti-CD147 antibody C (another two humanized anti-CD147 antibodies). The features of SARS-CoV-2, variants and SARS-CoV-2 pseudotyped variants and antibody inhibition rate were shown in Table 1. N501Y and E484K are mutation sites commonly found in B1.1.7, B1.1.351, B1.1.525, B1.1.526, P.1 and P.2 (Table 1). SPR analysis showed no significant difference in affinity constants between the RBD region of the spike protein and CD147 among the mutations (N501Y, E484K, Y453F and N439K) (Fig. 1F) and SARS-CoV-2 (*18*). These results indicate that SARS-CoV-2 and its variants can enter cells by CD147, and CD147 antibody can resist the infection of broad-spectrum variants.

**Fig. 1.**
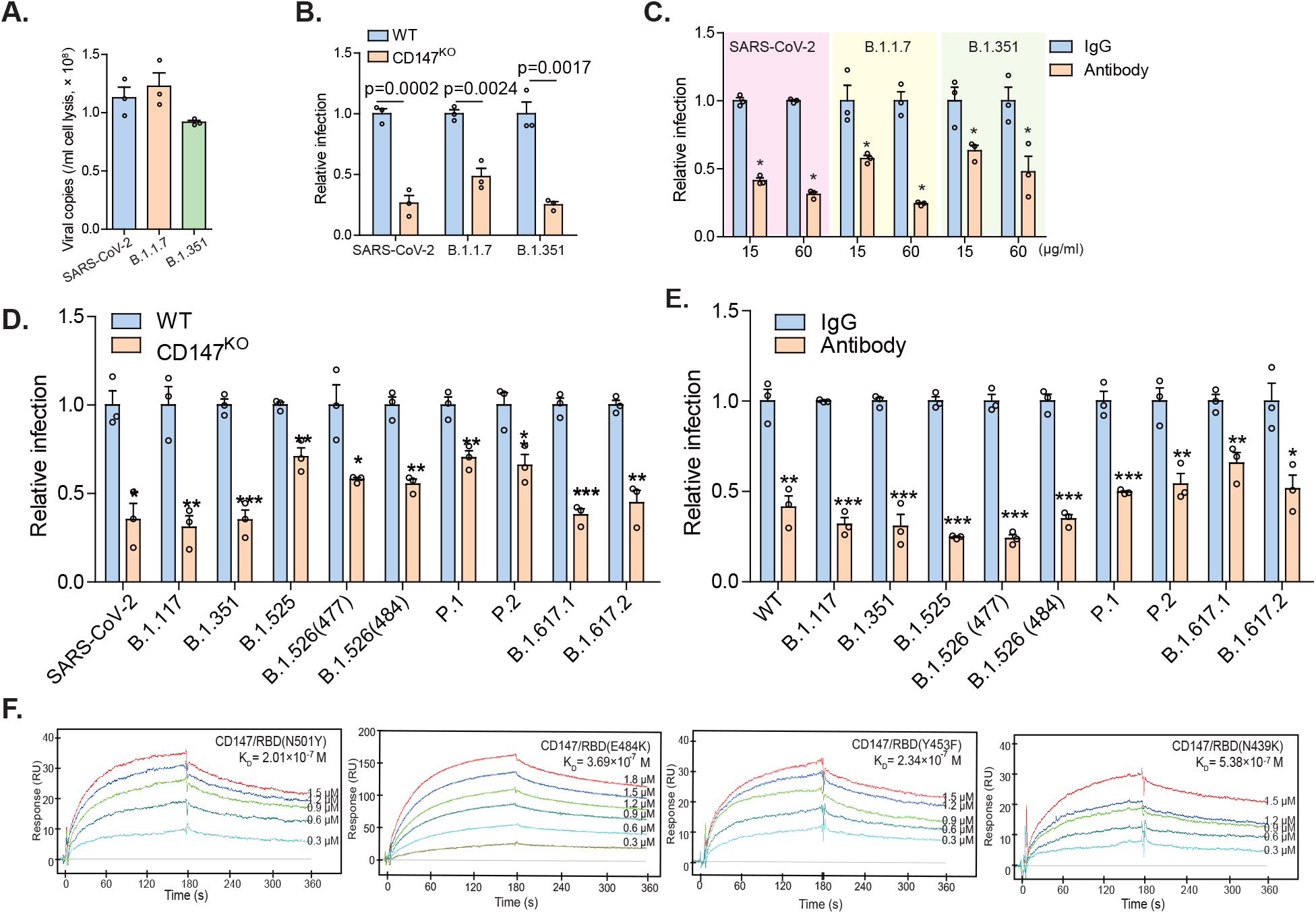
CD147 antibody effectively inhibits the infection of SARS-CoV-2 variants. SARS-CoV-2, B.1.1.7 and B.1.351 variants infected VeroE6 cells or CD147^KO^ VeroE6 cells for 48 h, and RNA was collected for viral RNA detection. **(A)** Taqman qRT-PCR for viral RNA level in SARS-CoV-2, B.1.1.7 or B.1.351 infected-VeroE6 cells. **(B)** Taqman qRT-PCR for viral RNA level in SARS-CoV-2, B.1.1.7 or B.1.351 infected-VeroE6 cells or CD147^KO^ VeroE6 cells. Data are normalized to the infectivity in wildtype VeroE6 cells (n = 3, two-tailed unpaired *t* test). **(C)** CD147 antibody inhibiting the infection of VeroE6 cells by SARS-CoV-2, B.1.1.7 and B.1.351 variants. Data are normalized to the infectivity in IgG group (n = 3, Two-tailed unpaired *t* test). **(D)** Viral infection efficiency detected by luciferase reporter assay. VeroE6 cells and CD147^KO^ VeroE6 cells were infected with the SARS-CoV-2 and 10 variants of SARS-CoV-2 pseudoviruses. Data are normalized to the infectivity in wildtype VeroE6 cells (n = 3, two-tailed unpaired *t* test). **(E)** CD147 antibody inhibiting the infection of VeroE6 cells by SARS-CoV-2 and 10 variants of SARS-CoV-2 pseudoviruses. B.1.1.7, B.1.351, B.1.525, B.1.526(S477N), B.1.526(E484K), P.1 and P.2 variants were inhibited by 60 μg/ml Meplazumab (a humanized anti-CD147 antibody), and B.1.617.1, and B.1.617.2 variants were inhibited by a combination of 30 μg/ml Mehozumab and 30 μg/ml anti-CD147 antibody C (another two humanized anti-CD147 antibodies). Data are normalized to the infectivity in IgG group (n = 3, Two-tailed unpaired *t* test). **(F)** The interaction of CD147 and four spike protein (RBD) mutants detected by SPR assay.

**Table 1.**
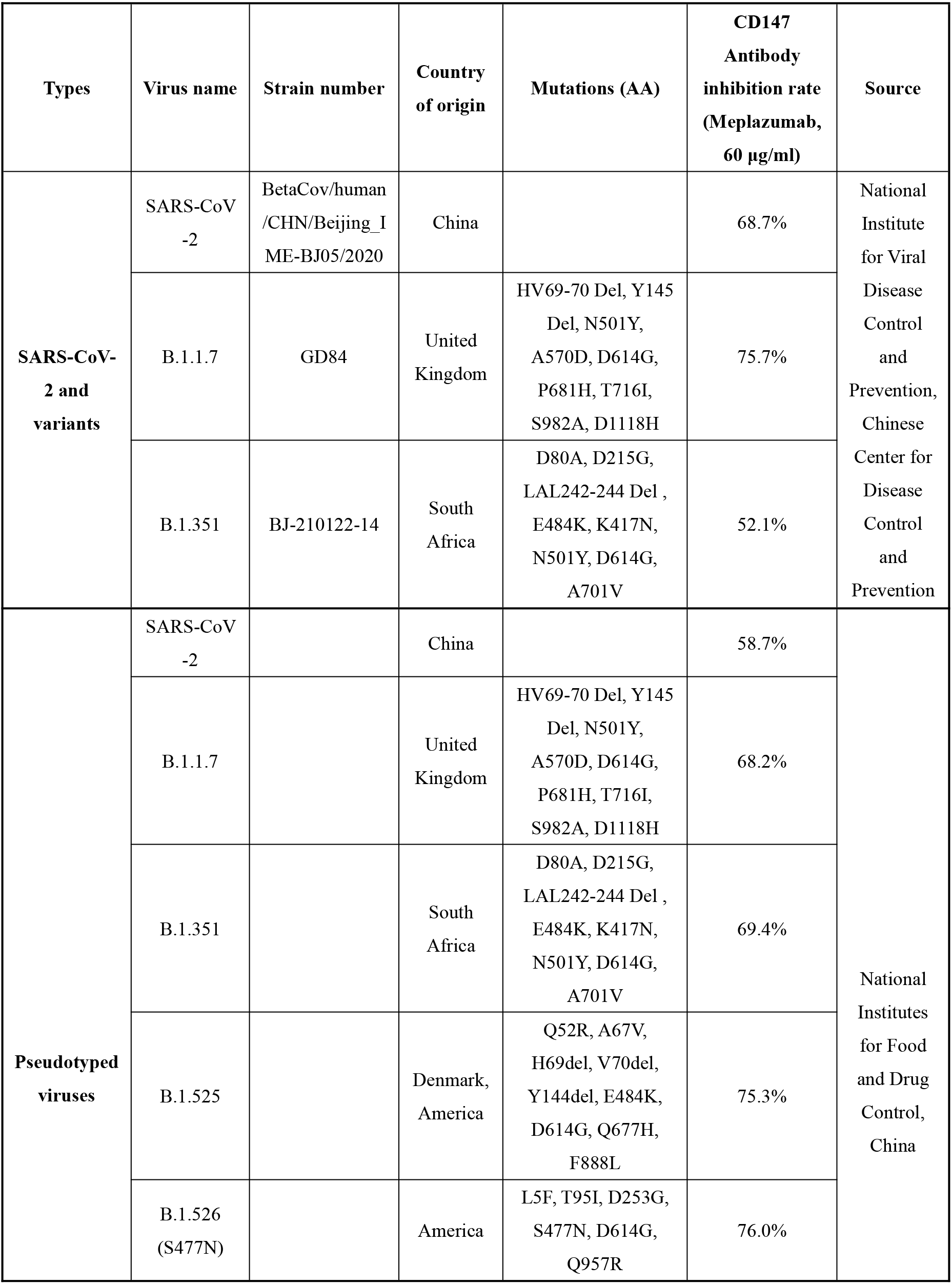
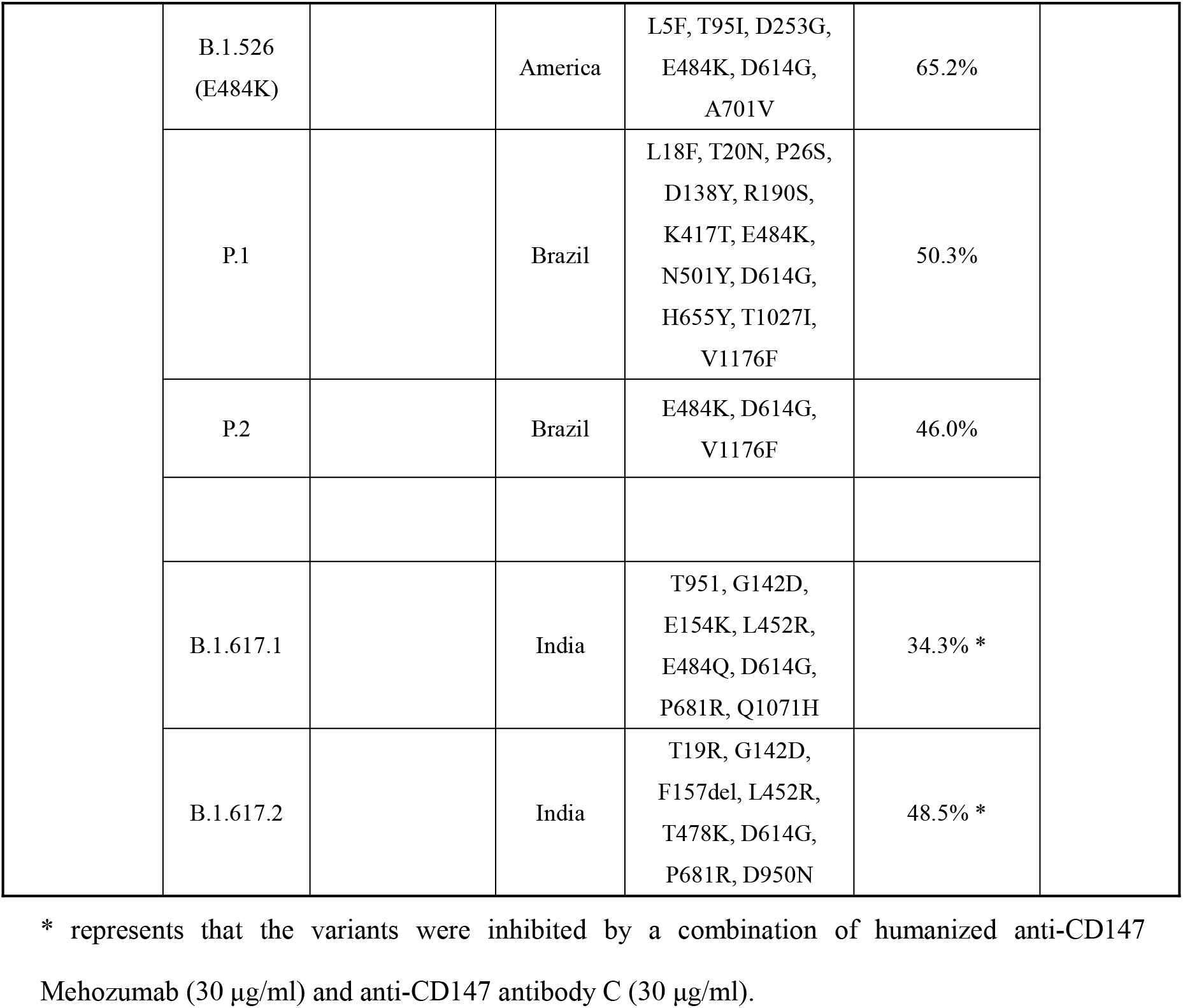
Features of SARS-CoV-2, variants and SARS-CoV-2 pseudotyped variants and CD147 antibody inhibition rate.

| Types | Virus name | Strain number | Country of origin | Mutations (AA) | CD147 Antibody inhibition rate (Meplazumab, 60 µg/ml) | Source |
| --- | --- | --- | --- | --- | --- | --- |
| **SARS-CoV-2 and variants** | SARS-CoV-2 | BetaCov/human /CHN/Beijing_I ME-BJ05/2020 | China |  | 68.7% | National Institute for Viral Disease Control and Prevention, Chinese Center for Disease Control and Prevention |
|  | B.1.1.7 | GD84 | United Kingdom | HV69-70 Del, Y145 Del, N501Y, A570D, D614G, P681H, T716I, S982A, D1118H | 75.7% |  |
|  | B.1.351 | BJ-210122-14 | South Africa | D80A, D215G, LAL242-244 Del , E484K, K417N, N501Y, D614G, A701V | 52.1% |  |
| **Pseudotyped viruses** | SARS-CoV-2 |  | China |  | 58.7% | National Institutes for Food and Drug Control, China |
|  | B.1.1.7 |  | United Kingdom | HV69-70 Del, Y145 Del, N501Y, A570D, D614G, P681H, T716I, S982A, D1118H | 68.2% |  |
|  | B.1.351 |  | South Africa | D80A, D215G, LAL242-244 Del , E484K, K417N, N501Y, D614G, A701V | 69.4% |  |
|  | B.1.525 |  | Denmark, America | Q52R, A67V, H69del, V70del, Y144del, E484K, D614G, Q677H, F888L | 75.3% |  |
|  | B.1.526 (S477N) |  | America | L5F, T95I, D253G, S477N, D614G, Q957R | 76.0% |  |

|  |  |  |  |  |  |
| --- | --- | --- | --- | --- | --- |
|  | B.1.526<br>(E484K) |  | America | L5F, T95I, D253G,<br>E484K, D614G,<br>A701V | 65.2% |
|  | P.1 |  | Brazil | L18F, T20N, P26S,<br>D138Y, R190S,<br>K417T, E484K,<br>N501Y, D614G,<br>H655Y, T1027I,<br>V1176F | 50.3% |
|  | P.2 |  | Brazil | E484K, D614G,<br>V1176F | 46.0% |
|  | B.1.617.1 |  | India | T95I, G142D,<br>E154K, L452R,<br>E484Q, D614G,<br>P681R, Q1071H | 34.3% * |
|  | B.1.617.2 |  | India | T19R, G142D,<br>F157del, L452R,<br>T478K, D614G,<br>P681R, D950N | 48.5% * |

\* represents that the variants were inhibited by a combination of humanized anti-CD147 Mehozumab (30 µg/ml) and anti-CD147 antibody C (30 µg/ml).

### The pathological and immune characteristics of hCD147 transgenic mice infected with SARS-CoV-2 are similar to those of COVID-19 patients

A variety of cytokines and chemokines were detected on VeroE6 cells infected by SARS-CoV-2, B.1.1.7 and B.1.351 variants, and it was found that the expressions of cytokines and chemokines caused by the three strains were comparable (Fig. S1). Therefore, SARS-CoV-2 was used as the main strain to detect the pathogenicity *in vivo*.

In order to detect the pathogenicity of SARS-CoV-2 via CD147, we constructed a human CD147 (hCD147) transgenic mouse model by replacing extracellular region of mouse CD147 with human CD147 in embryonic stem cells (Fig. S2A). PCR and FACS analyses showed that human CD147 was successfully expressed in mouse cells (Fig. S2B and C). We then inoculated hCD147 mice via the intranasal route with 3 × 10^5^ TCID_50_ of SARS-CoV-2. After 5 days post infection (d.p.i.), hCD147 mice demonstrated marked weight loss, and all mice lost approximately 10% of their body weight by 13 d.p.i. (Fig. 2A). High levels of viral RNA were detected in lung homogenates at 2 d.p.i. (Fig. 2B). In some of the mice, low levels of viral RNA were detected in organs, such as heart, spleen and kidney (Fig. 2B). H&E-stained lung sections from virus-infected hCD147 mice showed distinctive pathological characteristics at three time points, with the most severe symptoms at 6 d.p.i., characterized by widened alveolar wall, massive inflammatory cell infiltration, obvious alveolar serous fluid exudation and local lung consolidation (Fig. 2C and S2D). Multiplex immunofluorescence staining showed that a large number of macrophages, T cells and neutrophils infiltrated the lung alveoli and interstitium at 6 d.p.i. (Fig. 2D and E). In addition, infiltrated macrophages and neutrophils were also detected in the alveolar cavities (Fig. 2D). We found SARS-CoV-2 infection elevated several proinflammatory cytokines and chemokines, such as IL6, CXCL2, CXCL10, IFNγ and IL17a, in the lung tissues of hCD147 mice, compared with that of virus-infected C57BL/6J mice at 6 d.p.i. (Fig. 2F). RNA-seq analysis of lung homogenates from C57BL/6J and hCD147 mice at 2 d.p.i. showed 354 upregulated and 175 downregulated genes (Fig. S2E). KEGG analysis showed intensive inflammatory responses in hCD147 mice, including Th17 cell differentiation, Th1 and Th2 cells differentiation and MAPK pathway (Fig. 2G), similar to the immune responses in COVID-19 patients. The levels of cytokines and chemokines (IL6, IL10, IL1b, CXCL1, CXCL2 and CCL2) increased in the lung tissues of hCD147 mice, which was also found elevated in COVID-19 patients (*19–21*). In addition, inflammation-related transcription factors were analyzed increased in the lung tissues of virus-infected hCD147 mice (Fig. 2H). These results suggest that CD147 mediates potent inflammatory responses to SARS-CoV-2 infection, and the pathological and immune characteristics of hCD147 mice infected with SARS-CoV-2 imitate those of COVID-19 patients, which is an ideal model for study of pathogenesis of COVID-19.

**Fig. 2.**
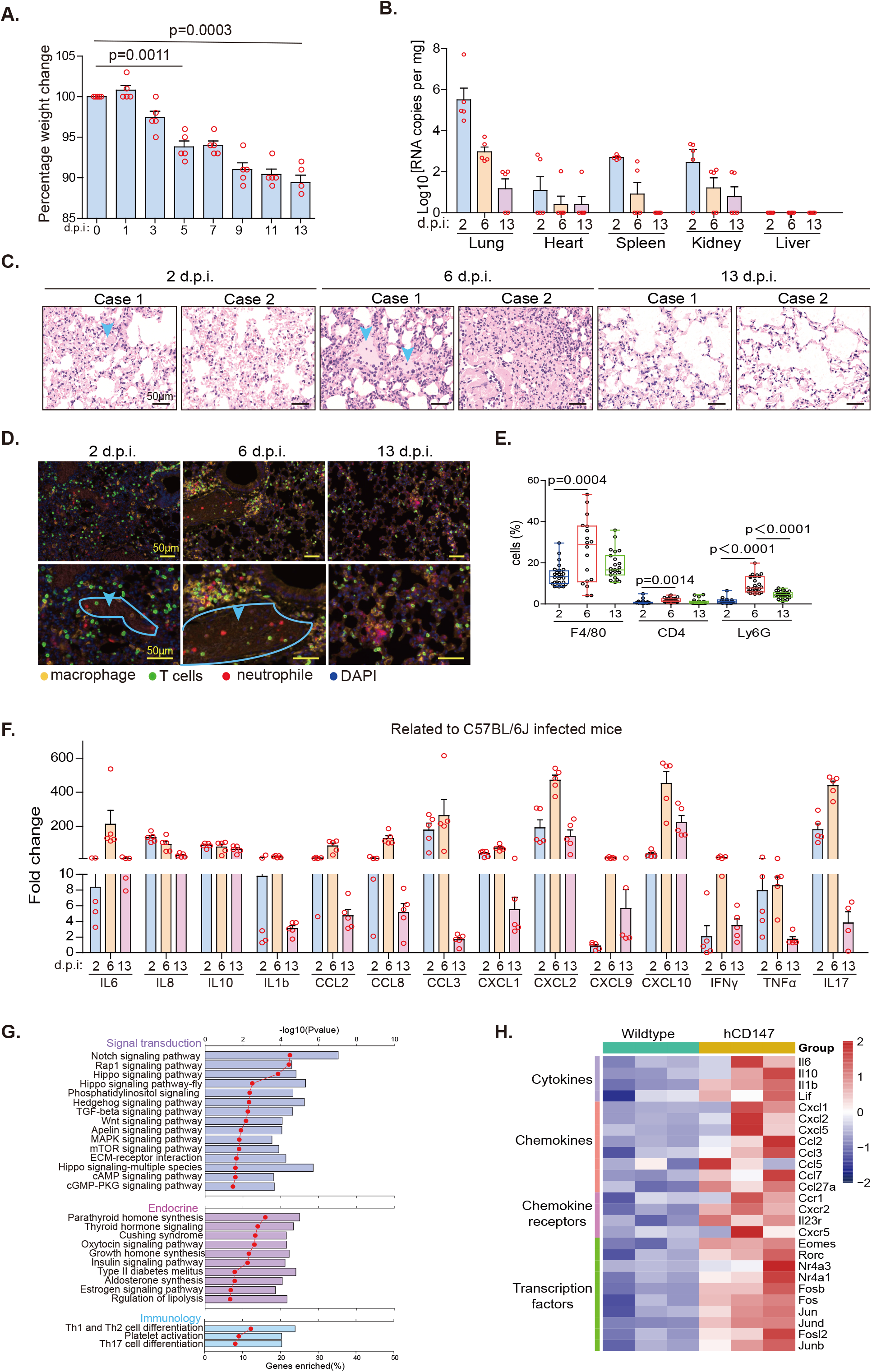
A hCD147 mouse model with susceptibility to SARS-CoV-2 infection and an inflammatory phenotype is established. hCD147 mice were inoculated via the intranasal route with 3×10^5^ TCID_50_ of SARS-CoV-2. **(A)** Weight changes were monitored (n = 5). **(B)** RT-qPCR for viral RNA levels in the lung, heart, spleen, kidney and liver (n = 5). **(C)** H&E staining of lung tissue sections from hCD147 mice at 2, 6 and 13 d.p.i. (scale bars, 50 μm). **(D)** Multiplex immunofluorescence staining of lung tissue sections from hCD147 mice at 2, 6 and 13 d.p.i. (scale bars, 50 μm). Cyan outline indicates exudation. **(E)** Statistics of the percentage of cells positive for CD3, F4/80 and Ly6G (n = 5, two-tailed unpaired *t* test). **(F)** Fold change in the gene expression of the cytokines and chemokines determined by RT-qPCR, compared with C57BL/6J controls in lung homogenates at 2, 6 and 13 d.p.i. Gapdh is used as a reference gene (n = 5). **(G)** RNA-seq analysis of the lung homogenates of C57BL/6J and hCD147 mice at 2 d.p.i. (n = 3). KEGG enrichment analysis of pathways enriched in changed genes. **(H)** Heat map of significantly upregulated genes during SARS-CoV-2 infection at 2 d.p.i. Columns represent samples and rows represent genes. Gene expression levels in the heat maps are z score–normalized values determined by log_2_^[CPM]^ values.

### SARS-CoV-2 causes different pneumonia phenotypes and immune responses in the hACE2 and hCD147 mouse models

To understand the pathological process caused by SARS-CoV-2 infection through different receptors, we infected hACE2 mice (*22*) and hCD147 mice with SARS-CoV-2. At 2 d.p.i. hACE2 mice presented interstitial pneumonia, whereas hCD147 mice showed exudative alveolar inflammation with more exudation and alveolar wall damage (Fig. 3A). Electron microscopy revealed infiltration of macrophages and neutrophils, and leakage of erythrocytes, in alveoli of hCD147 mice, whereas these cells were rarely observed in alveoli of hACE2 mice (Fig. 3B). Virions presented in the alveolar type II cells, vascular endothelial cells and macrophages in both mouse models (Fig. 3B). In addition, exfoliated alveolar type II epithelial cells were observed in the lung tissues of hCD147 mice (Fig. 3B). More infiltration of macrophages and neutrophils were found in the lung tissues of hCD147 mice than hACE2 mice (Fig. 3C). RNA-Seq analysis showed that the upregulated genes in SARS-CoV-2-infected hCD147 mice were enriched in many cytokines and chemokines, with IL-17 signaling related genes (Fig. 3D). These results indicate that SARS-CoV-2 causes different pneumonia phenotypes and immune responses in the hACE2 and hCD147 mouse models, and that the proinflammatory responses mediated by CD147 is stronger, which may trigger the occurrence of CSS.

**Fig. 3.**
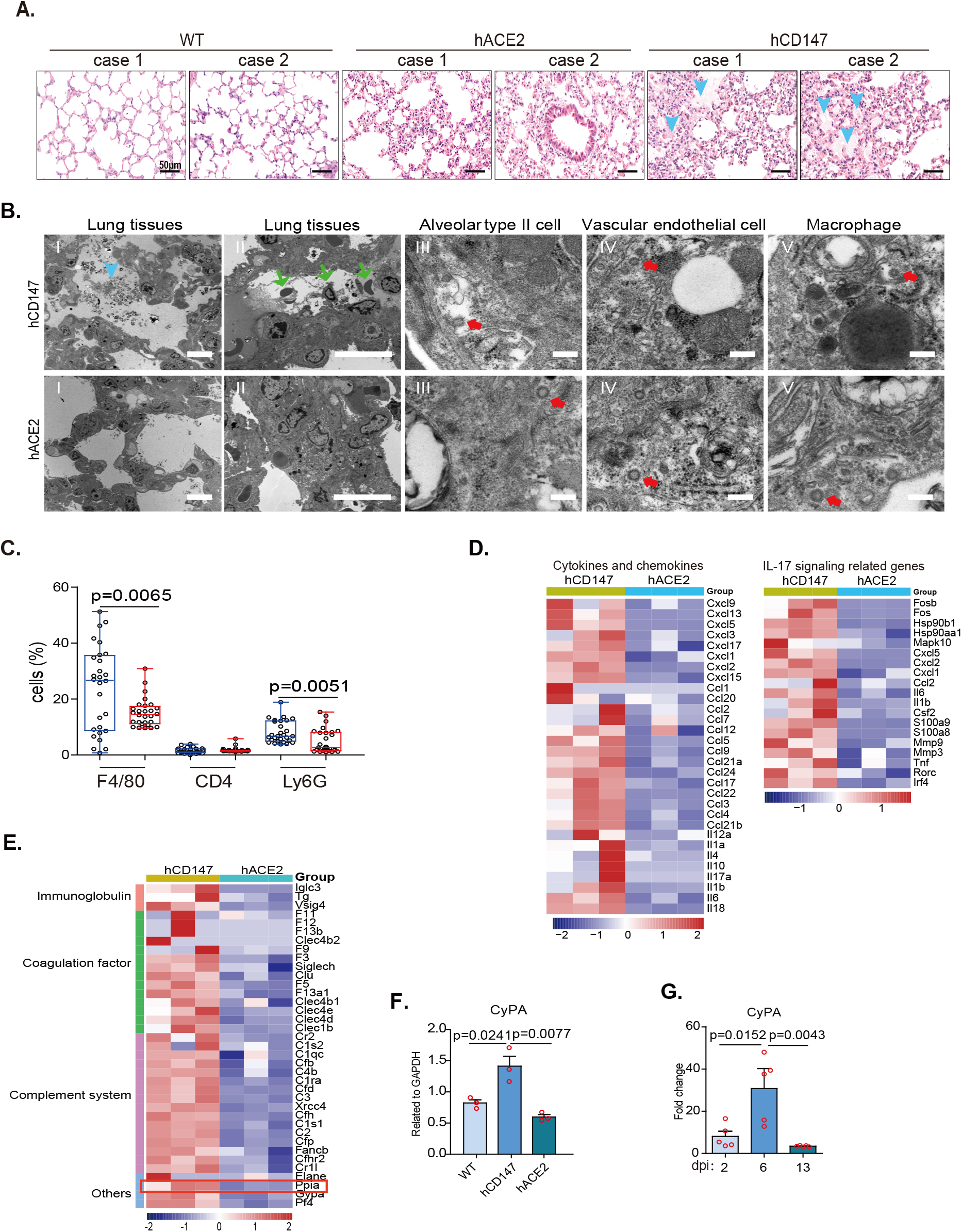
SARS-CoV-2 causes different pneumonia phenotypes and immune responses in the hACE2 and hCD147 mouse models. C57BL/6J, hACE2 and hCD147 mice were inoculated via the intranasal route with 3 × 10^5^ TCID_50_ of SARS-CoV-2. Lung tissues were collected at 2 d.p.i. for pathological analysis. **(A)** H&E staining of lung tissue sections from C57BL/6J, hACE2 and hCD147 mice (scale bars, 50 μm). **(B)** Pathological changes in hACE2 and hCD147 mouse lung tissues were detected by electron microscopy (scale bars in panels I and II, 10 μm; scale bars in panels III–V, 200 nm). The cyan arrow indicates exudation; the green arrows indicate infiltration of macrophages, neutrophils and erythrocytes; the red arrows indicate virions. **(C)** Statistics of the percentages of cells positive for CD3, F4/80 and Ly6G, stained by multiplex immunofluorescence (n = 6, two-tailed unpaired *t* test). **(D)** Heat map of significantly upregulated cytokines, chemokines IL-17 signaling related genes in lung homogenates of hCD147 mice versus hACE2 mice. Columns represent samples and rows represent genes. Gene expression levels in the heat maps are z score – normalized values determined by log2^[CPM]^ values. **(E)** Heat map of immune related genes (except cytokines and chemokines) in lung homogenates of hACE2 mice versus hCD147 mice. Columns represent samples and rows represent genes. Gene expression levels in the heat maps are z score–normalized values determined by log ^[CPM]^ values. **(F)** RT-qPCR for CyPA levels in lung homogenates of C57BL/6J, hACE2 and hCD147 mice (n = 3, two-tailed unpaired *t* test). **(G)** RT-qPCR for CyPA levels in lung homogenates of hCD147 mice after intranasal infection at 2, 6 and 13 d.p.i. (n = 5, two-tailed unpaired *t* test).

In order to make clear the molecular mechanism by which SARS-CoV-2 mediated different pathogeny through CD147 and ACE2 infection, we analyzed the upregulated immune related genes in hCD147 infected mice. In addition to cytokines and chemokines, the highly expressed genes related to the immune system were immunoglobulin, coagulation factor, complement system and so on (Fig. 3E). Notably, CyPA (PPIA), which is an upstream regulator of effectors such as IL6, IL1b, IL1a, CXCL1 and CXCL2 (*23*), showed significant upregulation in hCD147 mice (Fig. 3E). We verified CyPA expression in the lung tissues of infected mice by RT-qPCR and found that CyPA expression in hCD147 mice was considerably higher than that in hACE2 mice (Fig. 3F). Compared with C57BL/6J mice, CyPA expression in the lung tissues of virus-infected hCD147 mice was significantly increased at 2 d.p.i., peaked at 6 d.p.i. and reached a low level at 13 d.p.i. (Fig. 3G), which suggests the involvement of CyPA in inflammation caused by SARS-CoV-2 infection. Immunohistochemical staining further confirmed that the expression of IL6, CCL2 and CyPA was higher in the lung tissues of hCD147 mice than hACE2 mice (Fig. S3A, B and C). In addition, compared with hACE2 mice, total p-STAT3 expression was higher in lung tissues of hCD147 mice (Fig. S3D and E). The intensity of p-STAT3 was positively correlated with the CyPA expression in the CyPA-positive cells in hCD147 mice (Fig. S3F). These results suggest that CyPA is involved in the inflammatory responses of SARS-CoV-2 infection through CD147 and is likely to be a driver in the development of COVID-19 CSS.

### CyPA is a key intermediate proinflammatory factor involved in CD147-related immune responses upon SARS-CoV-2 infection

To further determine whether CyPA is a key intermediate proinflammatory factor that promotes the pathological process of SARS-CoV-2 infection via CD147, we conducted experiments in cell lines. In VeroE6 cells, upregulation of CyPA was induced by different titer virus infection (Fig. 4A). Although the virus content was reduced by CD147 or ACE2 knockdown (Fig. S4A), CyPA was detected downregulation by CD147 knockdown, but not by ACE2 knockdown (Fig. 4B). In addition, SARS-CoV-2 failed to infect VeroE6 cells with double knockout of CD147 and ACE2 (Fig. 4C), suggesting ACE2 and CD147 are the main receptors found for virus entry at present. CyPA expression decreased significantly in double knockout cells (Fig. 4D), and enhanced by CD147 overexpression (Fig. 4E). Previously, CyPA was reported to be regulated by the JAK-STAT pathway (*24*). Indeed, CyPA level could be inhibited by JAK inhibitor (Fig. 4E), indicating CD147 induces CyPA expression via JAK pathway during virus infection. Furthermore, the expression of CD147 and CypA was measured by multiplex immunofluoresence staining in lung tissues from COVID-19 patients (Fig. S4B), and the results showed that CyPA expression was significantly elevated in CD147^high^ regions than CD147^low^ regions (Fig. S4C). The expression of CD147 had a strong positive correlation with CyPA (Fig. S4D), which further verified that CD147 upregulated CyPA expression upon SARS-CoV-2 infection.

**Fig. 4.**
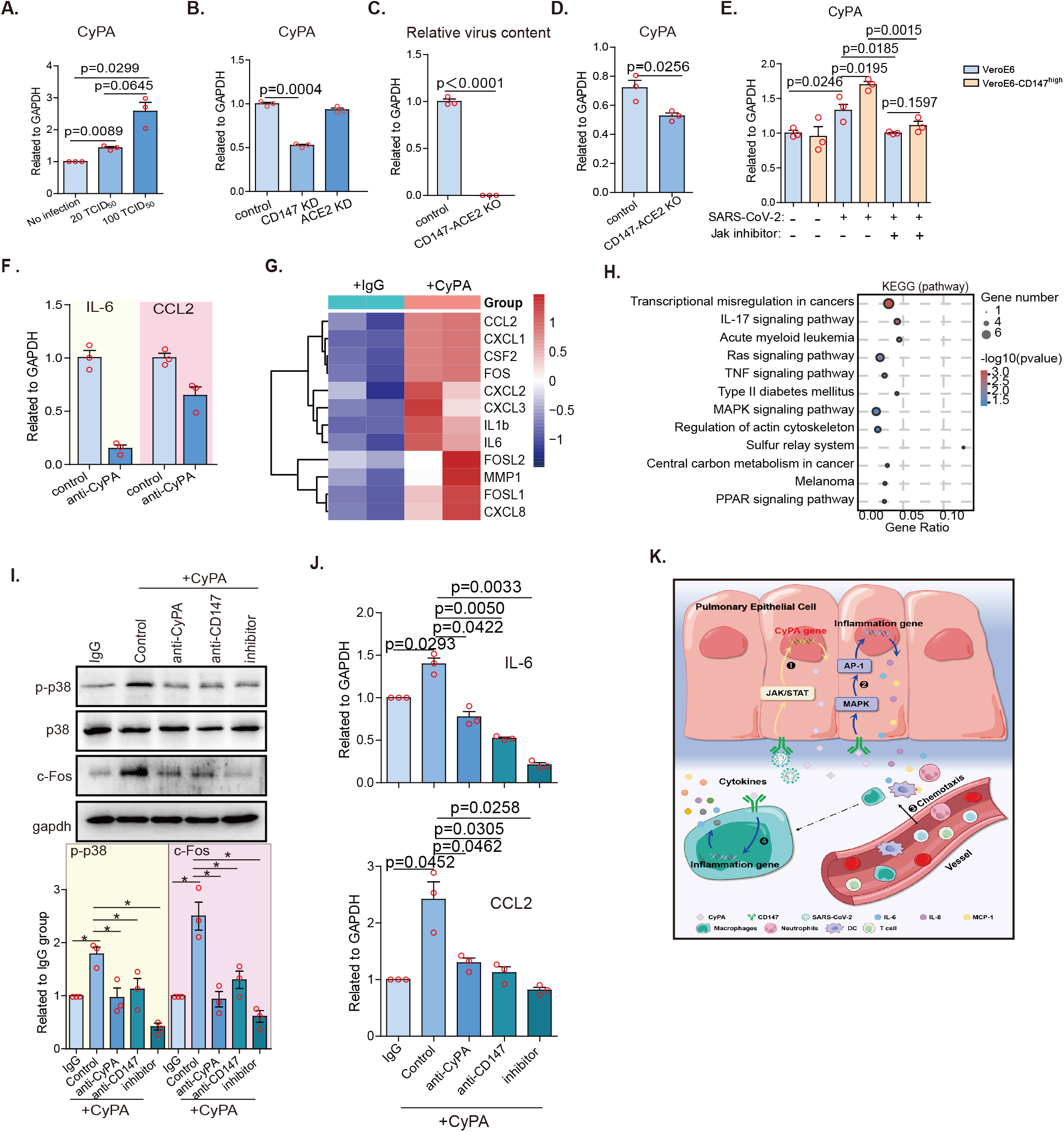
CyPA is a key intermediate proinflammatory factor involved in CD147-related immune responses. **(A)** SARS-CoV-2-infected VeroE6 cells with different titers were incubated for 48 h, and RNA was collected for CyPA detection by RT-qPCR (n = 3, two-tailed unpaired *t* test). **(B)** CD147 or ACE2 in VeroE6 cells was knocked down by lentivirus, and the cells were then infected with SARS-CoV-2 for 48 h. The expression level of CyPA was detected by RT-qPCR (n = 3, two-tailed unpaired *t* test). **(C)** CD147 and ACE2 in VeroE6 cells were both knocked out by CRISPR-Cas9, the cells were then infected with SARS-CoV-2 for 48 h. Viral RNA was detected by RT-qPCR, normalized to Gapdh (n = 3, two-tailed unpaired *t* test). **(D)** CyPA expression was detected by RT-qPCR in double knockout of CD147 and ACE2 VeroE6 cells infected with SARS-CoV-2 for 48 h, normalized to Gapdh (n = 3, Two-tailed unpaired *t* test). **(E)** CD147 in VeroE6 cells was ectopically expressed by lentivirus, the cells were then infected with SARS-CoV-2 for 48 h with or without JAK inhibitor. The expression level of CyPA was detected by RT-qPCR (n = 3, two-tailed unpaired *t* test). **(F)** IL6 and CCL2 expression was detected by RT-qPCR in VeroE6 cells infected with SARS-CoV-2 for 48 h with or without CyPA antibody, compared with the control (n = 3, two-tailed unpaired *t* test). Gapdh is used as a reference gene. **(G)** Heat map of significantly upregulated genes based on RNA-seq. BEAS-2B cells were stimulated with or without CyPA for 48 h. Columns represent samples and rows represent genes. Gene expression levels in the heat maps are z score–normalized values determined by log_2_^[CPM]^ values. **(H)** KEGG enrichment analysis of pathways enriched in upregulated genes from the CyPA-stimulated group compared with the control group. **(I)** Western blot analysis of p-p38, p-38 and c-FOS expression (n = 3, two-tailed unpaired *t* test, *p < 0.05). **(J)** IL6 and CCL2 expression detected by RT-qPCR. **(K)** Schematic diagram of CD147-CyPA regulation of cytokine expression.

CyPA is an intracellular protein that can also be secreted extracellular, and its most advantageous place to exert its role as a driver of COVID-19 CSS is extracellular. Therefore, CyPA antibody was used to block the extracellular CyPA to evaluate whether the expression of cytokines and chemokines would be affected. Evidently, the anti-CyPA antibody reduced the expression of IL6 and CCL2 (Fig. 4F), with the viral load unchanged (Fig. S4E). To map the CyPA-related transcriptome networks, RNA-seq was performed using human lung epithelial BEAS-2B cells, that were incubated with CyPA. In consistent with the *in vivo* data from SARS-CoV-2-infected hCD147 mice, CyPA-stimulated BEAS-2B cells showed upregulated expression of a variety of proinflammatory cytokines and chemokines, including IL6, IL1b, CCL2, CXCL1 and CXCL2, as well as several AP-1 family members, including c-FOS, FOSL1 and FOSL2 (Fig. 4G) that promote the transcription of multiple cytokine genes during inflammation (*25*). Kyoto Encyclopedia of Genes and Genomes (KEGG) analysis showed significant enrichment in the MAPK pathway (Fig. 4H) which regulates AP-1 (*26*), suggesting that CyPA may regulate cytokine expression through activating the MAPK-AP-1 pathway. CD147 is a major receptor for extracellular CyPA (*27*), with an affinity constant K_D_ of 3.34 × 10^−8^ M (Fig. S4F and G). Western blot analysis showed that CyPA induced an increase in p-p38 and c-FOS expression, which could be reversed by either anti-CyPA antibody, anti-CD147 antibody or MAPK inhibitor (Fig. 4I). In addition, the expression of IL6 and CCL2 was effectively inhibited by the above three agents (Fig. 4J). These results indicate that CyPA induces cytokine expression through the CD147-mediated MAPK pathway (Fig. 4K).

### CyPA is involved in inflammatory responses in COVID-19 patients

To confirm the clinical significance of CyPA in the inflammatory responses during COVID-19, a cytokine chip assay was performed and showed that 32 out of 40 cytokines were upregulated in plasma of COVID-19 patients (n = 200, including 41 severe/critical and 159 mild), compared with healthy individuals (n = 100) (Fig. 5A and S5). Meanwhile, CyPA level in plasma analyzed by ELISA was significantly higher in severe/critical COVID-19 patients (n = 106, comprising the above 41 severe/critical cases) than in mild (n = 159) and healthy cohorts (n = 100) (Fig. 5B). Further, CyPA level in clinical samples was significantly correlated with those above upregulated cytokine level, especially IL17, CCL2 and IL6 (Fig. 5C).

**Fig. 5.**
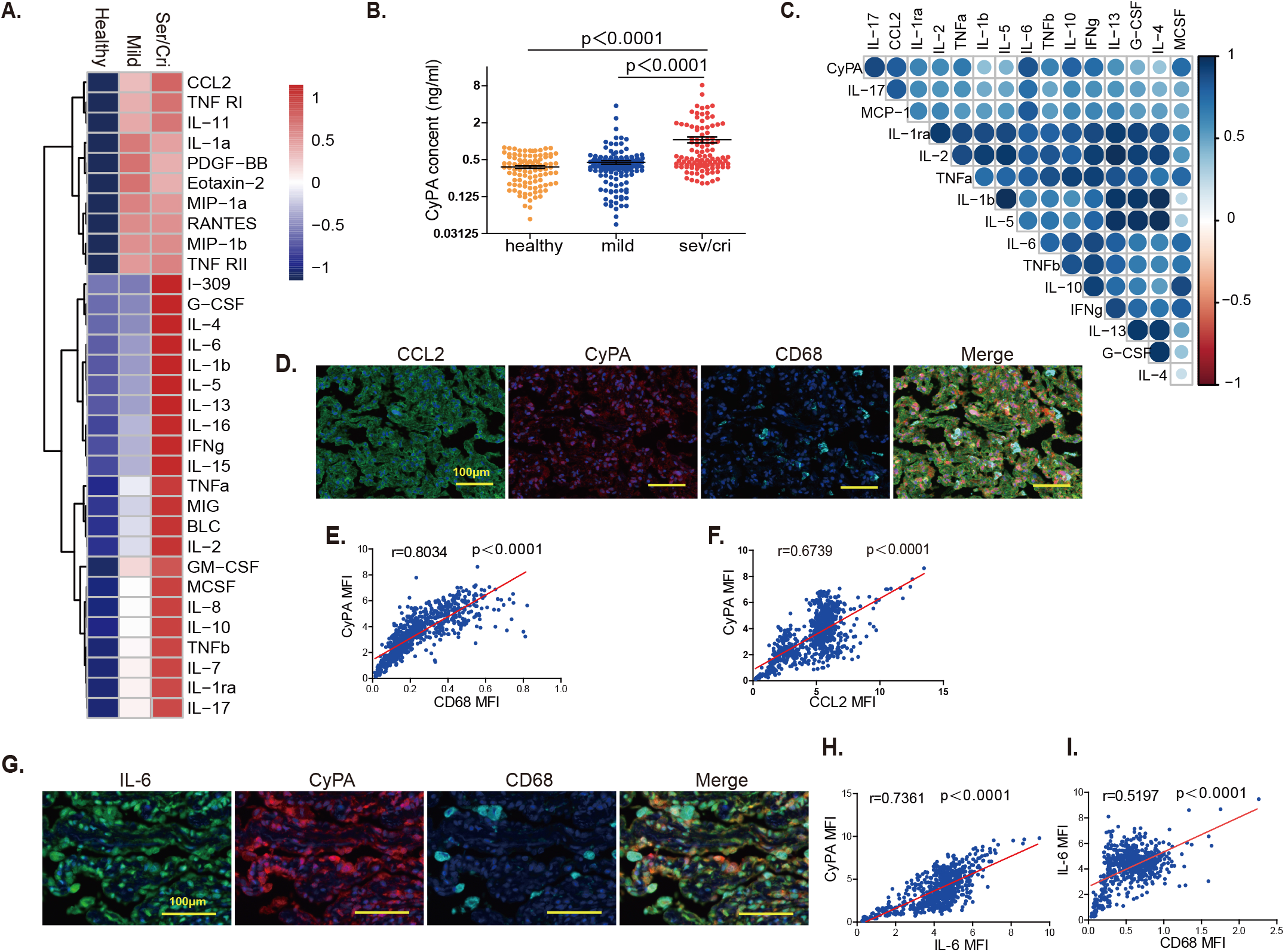
CyPA is involved in inflammatory responses in COVID-19 patients. **(A)** The heat map of 32 differential cytokines. 40 human cytokines were detected in plasma from healthy individuals (n = 100) and COVID-19 patients (n = 200, including 41 severe/critical and 159 mild) using cytokine chips. **(B)** CyPA protein detected in the plasma of healthy donors (n = 100), COVID-19 patients with mild symptoms (n = 159) and COVID-19 patients with severe/critical symptoms (n = 106) by ELISA (two-tailed unpaired *t* test). **(C)** Correlation analysis between CyPA and cytokines, according to the Pearson correlation coefficient. Pathological analysis of lung tissues from 13 cases COVID-19 patients. **(D)** Multiplex immunofluorescence staining was carried out for CyPA, CD68 and CCL2 (scale bars, 100μm). **(E)** Correlation analysis of CyPA and CD68. **(F)** Correlation analysis of CyPA and CCL2. **(G)** Multiplex immunofluorescence staining was carried out for CyPA, CD68 and IL6 (scale bars, 100μm). **(H)** Correlation analysis of CyPA and IL6. **(I)** Correlation analysis of IL6 and CD68. All the correlation scatter plots represent 13 cases of COVID-19 lung tissues data, which were analyzed by HALO software. All correlation analysis performed by Pearson correlation.

We then collected lung tissues from 13 cases of COVID-19 patients through puncture or autopsy and stained them with H&E staining (Fig. S6A). In order to further clarify the correlation between CyPA and infiltrating cells and cytokines, multiplex immunofluorescence staining was carried out to detect the expression of CyPA, macrophage marker CD68, CCL2 and IL6 (Fig. 5D and G). Each collected image was divided into nine process regions, and the MFI of each marker in each region was analyzed with Halo software and correlation analysis was performed (Fig. S6B). The results showed that in the lung tissues of patients with COVID-19, CyPA had a strong positive correlation with macrophages and CCL2 (Fig. 5E and F). In particular, the correlation coefficient reached 0.8 between CyPA and macrophages, indicating that CyPA is related to the infiltration of macrophages. In addition, most of cells (including macrophages) in the lung tissues of COVID-19 strongly expressed IL6 (Fig. 5G), which had a positive correlation with CyPA (Fig. 5H), suggesting that CyPA promotes IL6 expression. These results indicate that CyPA is involved in inflammatory responses in COVID-19. In addition, there was a correlation between IL6 and macrophages, but the correlation coefficient was only 0.5 (Fig. 5I), further indicating that not only the macrophages secrete IL6, but other cells in the lung tissues of COVID-19 could also highly express IL6.

### CD147 antibody reverses pulmonary inflammation caused by SARS-CoV-2

To verify whether CD147 functions as a therapeutic target for COVID-19, we treated SARS-CoV-2-infected hCD147 mice with CD147 antibody (Meplazumab) (Fig. S7A). CD147 antibody effectively cleared the virus and alveolar exudation, resolving the pneumonia at 6 d.p.i. (Fig. 6A, B and S7B). CD147 antibody treatment had a favorable therapeutic effect on the downregulation of cytokines and chemokines at 6 d.p.i., such as IL6, IL10, IL1b, CCL2, CXCL1, CXCL2, IFN-γ, IL-17 and CyPA (Fig. 6C). Infiltration of macrophages and neutrophils was decreased after the CD147 antibody treatment (Fig. 6D). These results indicate that CD147 antibody is effective against COVID-19, supportive of our exploratory phase II clinical trial results (*28*). Therefore, CD147 can be a promising target for the treatment of COVID-19 patients, especially those with severe pneumonia.

**Fig. 6.**
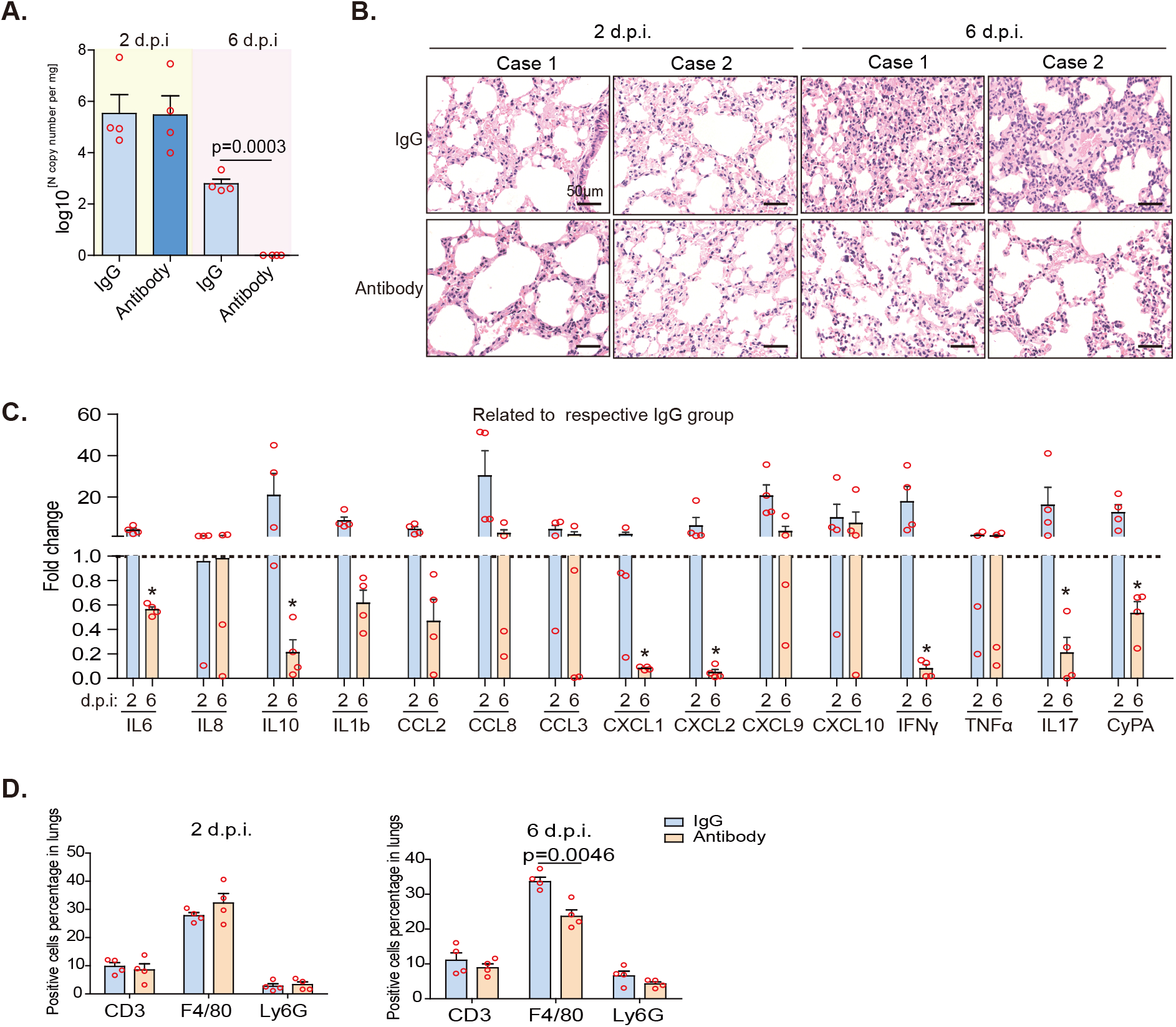
CD147 antibody reverses pulmonary inflammation caused by SARS-CoV-2. hCD147 mice were intranasally inoculated with 3 × 10^5^ TCID_50_ SARS-CoV-2 and treated with CD147 antibody Meplazumab the next day. Lung tissues were collected at 2 and 6 d.p.i. **(A)** RT-qPCR for viral RNA levels in lung tissues (n = 4). **(B)** H&E staining of lung tissue sections from the IgG and CD147 antibody groups at 2 and 6 d.p.i. (scale bars, 50 μm). **(C)** Fold change in the gene expression of cytokines and chemokines in lung homogenates determined by RT-qPCR, compared with the corresponding IgG controls at 2 and 6 d.p.i. (n = 4, two-tailed unpaired *t* test, *p < 0.05). Gapdh is used as a reference gene. **(D)** Statistics of the percentages of cells positive for CD3, F4/80 and Ly6G at 2 and 6 d.p.i. (n = 4, two-tailed unpaired *t* test).

## Discussion

SARS-CoV-2 variants are raising global concerns not only because of their increased transmission but also because of their extensive mutations in spike protein that could lead to antigenic changes detrimental to neutralization of antibody therapy and vaccine protection. D614G, occurred at the initial stage of the pandemic, is the most common mutant, whichmakes the virus more infectious and transmissible (*29*). N501Y, commonly appeared in B.1.1.7, B.1.351 and P.1 variants, allows the virus to more readily bind to ACE2 (*30*). E484K, first found in B.1.351 variant, helps the virus slip past the body’s immune defense (*31*). The uncertainty of spike protein mutation considerably undermines the efficacy of therapeutics and vaccination, thus calling for new strategies against SARS-COV-2. Previous studies in our lab have found that the RBD of spike can bind with CD147 to help SARS-CoV-2 enter host cells (*18*). Our study showed that the four mutations located in the RBD domain, including N501Y, E484K, Y453F and N439K, do not change the affinity of RBD to CD147, indicating that CD147 still served as a receptor for SARS-CoV-2 variants. More importantly, we confirmed that CD147 antibody can effectively block the infection of the prevalent variants, such as B.1.1.7, B.1.351, P.1, B.1.526 and B.1.617, at an inhibition rate comparable to that of SARS-CoV-2. As such, CD147 is a shared receptor for SARS-CoV-2 and its variants, and CD147 antibody is a specific and effective receptor antagonist against COVID-19.

At present, another constraint for COVID-19 control is the lack of sufficient understanding of its pathogenesis. We investigated the pathogenesis of SARS-CoV-2 using hCD147 transgenic mice. Consistent with reports by others (*14, 32*), we observed interstitial pneumonia in virus-infected hACE2 mice, while hCD147 mice exhibited exudative alveolar pneumonia after SARS-CoV-2 infection. Moreover, hCD147 mice presented with Th17 cell response and stronger proinflammatory responses, which are similar to the initial immune responses of COVID-19 patients, contributing to the development of exudative alveolar pneumonia (*33–35*). Using this model, CD147 antibody effectively blocked the infection and inflammation induced by SARS-CoV-2, further verifying the effectiveness of CD147 antibody in the treatment of COVID-19. These findings also duplicated the results of the exploratory clinical trial of CD147 antibody against SARS-CoV-2, which showed favorable outcomes in discharge rates, recovery rates of critical severe patients, lung inflammation and time to clearance of virus (*28*). Thus, SARS-CoV-2-infected hCD147 transgenic mice well recapitulate the symptoms of COVID-19 pneumonia, providing a good model for studying the pathogenic mechanism and pharmacology of agents against SARS-CoV-2.

Cytokine storm caused by SARS-CoV-2 is a main cause of death in severe COVID-19 patients, and Th17 cell response contributes to the cytokine storm during viral infection, which results in tissue damage and pulmonary edema (*35*). The infection of SARS-CoV-2 via CD147 can induce Th17 cell response, suggesting that CD147 is the key factor to cytokine storm. In this process, CyPA is an important intermediate product, which is considered to be a critical proinflammatory factor (*36*). The spike protein of SARS-CoV-2 binding to CD147 activates JAK-STAT3 pathway, thus promoting the expression of CyPA. A high level of CyPA not only induces chemotaxis in numerous monocytes and neutrophils to the inflammatory site, but also activates the MAPK signaling pathway by binding with CD147 to promote the massive production of cytokines, such as CCL2 and IL6. Hence, in the process of SARS-CoV-2 infection, CyPA is a key mediator that enhances the inflammatory response, and CD147, as a regulatory molecule of JAK-STAT3 upstream, triggers the cytokine storm.

Taken together, CD147 plays a central role in SARS-CoV-2 and its variants infection and pathogenesis of COVID-19. CD147 is not only a viral entry receptor, but also a signal initiator for cytokine storm. Accordingly, blocking CD147 inhibits SARS-CoV-2 infection and suppresses cytokine storm. In conclusion, CD147 is a core target for SARS-CoV-2 variants and CD147 antibody is a specific and effective drug to control ever-spreading COVID-19 epidemic.

## Acknowledgments

We thank Prof. Chuan Qin from the Chinese Academy of Medical Sciences for helping to conduct the animal experiments.

## Author contributions

Experimental design: C.Z.N., Z.P., B.H.J., C.L. and G.J.J.

Experimental conduction: G.J.J., L.C., Y.Y.F., C.R., W.K., D.Y.Q., D.P., W.G.Z., W.Y.C., X.K., L.J.N., S.X.X., Y.X., W.J., J.J.L., L.L., Z.H., Z.K., Z.C.H., F.X.H., Y.F.F., Z.Z., W.D., Z.Y., G.T., S.Y., Z.Y.M., P.Z., H.F., C.S.R. and performed most of the experiments, including SPR, Co-IP, ELISA, PCR, SARS-CoV-2 strain isolation and TEM.

OpNS-EM experiment and image analysis: Y.X., C.S.R. and Z.Y. Mouse infection experiments: B.L.L, Z.H. L.J.N. and Q.C.

Provided and analysed the tissues from COVID-19 patients: Y.Y.F., X.W. and L.Y.W.

Data analysis: G.J.J., C.L., and C.R.

Supervision: G.J.J., C.L., Y.Y.F., C.R., W.K., and D.Y.Q.

Writing – original draft: G.J.J., B.H.J., and C.L.

Writing – review & editing: W.Q.Y., G.J.J., B.H.J., C.L., Z.P., and C.Z.N.

## RESOURCE AVAILABILITY

### Lead Contact

Further information and requests for resources and reagents should be directed to and will be fulfilled by the Lead Contact, Zhi-Nan Chen.

### Materials Availability

All materials and reagents generated in this study are available from the Lead Contact with a completed Materials Transfer Agreement.

### Data and Code Availability

The data that support the findings of this study are available from the corresponding author upon request, and has been uploaded to GEO (GSE167403).

## EXPERIMENTAL MODEL AND SUBJECT DETAILS

### Virus and cell lines

The SARS-CoV-2 strain (BetaCov/human/CHN/Beijing_IME-BJ05/2020) used for in vitro experiments was obtained from the State Key Laboratory of Pathogen and Biosecurity at Beijing Institute of Microbiology and Epidemiology. The SARS-CoV-2 strain (BetaCoV/Wuhan/IVDC-HB-01/2020/EPI_ISL_402119) used for in vivo experiments was from the Chinese Academy of Medical Sciences.

The VeroE6 and BEAS-2B cell lines were obtained from the Cell Bank of the Chinese Academy of Sciences (Shanghai, China). All cell lines were authenticated using short tandem repeat DNA profiling at Beijing Microread Genetics Co., Ltd. (Beijing, China) and cultured at 37°C under 5% CO2 in Dulbecco’s modified Eagle’s medium (DMEM, Invitrogen) or RPMI 1640 medium supplemented with 10% foetal bovine serum (FBS, Life Technologies), 1% penicillin/streptomycin and 2% L-glutamine.

### hACE2 and hCD147 mice

Human CD147 transgenic mice (hCD147) were provided by the Shanghai Model Organisms Center, Inc. (China). The extracellular region of CD147 in wild-type (WT) C57BL/6J mice was replaced with human CD147 by targeting embryonic stem cells. Human ACE2 transgenic mice (hACE2) were provided by the National Institutes for Food and Drug Control. These mice were bred and maintained in a specific pathogen-free facility at the Chinese Academy of Medical Sciences. The animal experiments were performed in accordance with the People’s Republic of China Legislation Regarding the Use and Care of Laboratory Animals. All protocols used in this study were approved by the Institutional Animal Care and Use Committee of Fourth Military Medical University (20200206).

### Plasma and lung tissue sections from COVID-19 patients

Plasma was obtained from healthy individuals (n = 100) at Xijing Hospital of the Fourth Military Medical University, and plasma was obtained from COVID-19 patients (n = 206) at Zhongnan Hospital of Wuhan University. Lung tissue sections were obtained from COVID-19 patients at Tangdu Hospital of the Fourth Military Medical University (n = 2) and the Institute of Clinical Pathology of the Third Military Medical University (n = 11).

This study was approved by the ethics committees of Xijing Hospital and Tangdu Hospital, Fourth Military Medical University (KY20202005-1, K202002-01, E202003-01), Third Military Medical University (2020074) and Zhongnan Hospital of Wuhan University (2020056K).

## METHOD DETAILS

### Generation of stable knockdown and knockout cell lines

VeroE6 cells were transfected with supernatants containing lentiviruses encoding the shCD147 and shACE2 constructs (GeneChem Co. Ltd.) using Lipofectamine 2000 reagent (Invitrogen) to generate the CD147 and ACE2 knockdown cell lines, respectively. After 48 h, the infected cells were selected with 3 μg/ml puromycin, and monoclonal cells were selected for further study. CD147-/-ACE2-/-VeroE6 cells were generated using the CRISPR/Cas9 system (GeneChem Co. Ltd).

### Infection of hCD147 and hACE2 mice with SARS-CoV-2

After being anaesthetized, each mouse was intranasally inoculated with SARS-CoV-2 at a TCID_50_ of 3 × 10^5^. For CD147 antibody treatment, 3 mg/kg CD147 antibody (Meplazumab) was administered via the tail vein at 1 d.p.i. Lung tissues were collected for RNA extraction or fixed with paraformaldehyde and PLP Fixing Solution for H&E staining, IHC, immunofluorescence staining and TEM.

### Histological analysis

Mouse lung samples were fixed with 4% paraformaldehyde, embedded in paraffin and cut into 3-μm-thick sections. Formalin-fixed paraffin-embedded mouse lung tissue sections were deparaffinized with xylene and alcohol. Fixed tissue samples were used for haematoxylin-eosin (H&E) staining and indirect IHC. For H&E staining, the slides were counterstained with haematoxylin for 15 min and eosin for 10 min. For IHC, the slides were deparaffinized and rehydrated, followed by 15 min of blocking with 5% goat serum in phosphate-buffered saline (PBS) and staining with primary antibodies for 2 h. Immunoperoxidase staining was performed using a General SP kit (SP-9000, ZSGB-BIO), and the sections were then treated with 3,3’-diaminobenzidine (ZLI-9019, ZSGB-BIO) to detect the target proteins, followed by counterstaining of nuclei with haematoxylin.

### Multiplex immunofluorescence staining

Multiplex immunofluorescence analyses were performed using 3-μm-thick sections of formalin-fixed and paraffin-embedded tissues. Briefly, slides were deparaffinized in xylene and hydrated in ethanol at gradient concentrations. After heat-induced antigen retrieval in either citrate (pH = 6) or Tris-EDTA (pH = 9) buffer, the samples were permeabilized with 0.5% Triton X-100, blocked with 5% goat serum-PBS and stained with primary antibodies. Nuclei were stained with DAPI. A TSA-indirect kit was used according to the manufacturer’s instructions (PerkinElmer). Image analysis was performed using HALO software (Akoya Biosciences).

### Quantitative measurement of cytokines

Forty human cytokines were detected by Quantibody® array kits (QAH-INF-3, RayBiotech) according to the manufacturer’s protocol. Then, 100 μl of standard cytokines or samples was added to each well after blocking, followed by incubation with the arrays at room temperature for 2 h. The wells were washed, and 80 μl of the detection antibody cocktail was added to each well and incubated at room temperature for 1 h. Eighty microlitres of Cy3 equivalent dye-conjugated streptavidin was added to each well and incubated in the dark at room temperature for 1 h. The signals were visualized by Axon GenePix. The data were analysed by microarray analysis software (GenePix, ScanArray Express, ArrayVision and MicroVigene).

### Enzyme-linked immunosorbent assay (ELISA)

The content of CyPA in plasma was detected using ELISA kits according to the manufacturer’s instructions (Eiaab Science, Inc., 0979 h). In brief, the microtiter plate provided in the kit was precoated with a biotin-conjugated antibody specific to the target CyPA. Standards were established at 6 different dilutions ranging from 0 ng/ml to 10 ng/ml, and nonimmune serum was used as a negative control. Next, 100 μl of the plasma, standard or negative control was added to each well of the plate and incubated for 2 h at 37°C. Then, avidin conjugated to horseradish peroxidase (HRP) was added to each well and incubated. The reaction was developed by adding 100 μl of 3,3’,5,5’-tetramethylbenzidine (TMB) to the wells for 12 min at room temperature. Finally, 200 M sulfuric acid was added to stop the reaction, and the absorbance at 450 nm was determined using a BioTeck Epoch microplate reader.

### RNA-seq

RNA was isolated using MyOne Silane Dynabeads (Thermo Fisher Scientific, Inc.). RNA was fragmented and barcoded using 8-base pair barcodes together with standard Illumina adaptors. The primers were removed using Agencourt AMPure XP (Beckman Coulter/Agencourt), and the samples were amplified by 14 PCR cycles. The libraries were gel purified and quantified using a Qubit high-sensitivity DNA kit (Invitrogen), and the library quality was confirmed using Tapestation High-Sensitivity DNA tapes (Agilent Technologies). The RNA-seq reactions were sequenced (50-base pair reads) on an Illumina NextSeq platform (Illumina) according to the manufacturer’s instructions, and analysis was performed using the CLC Genomics Workbench v.8.0.1 RNA-seq analysis software package (Qiagen). Briefly, the reads were aligned (mismatch cost = 2, insertion cost = 3, deletion cost = 3, length fraction = 0.8, similarity fraction = 0.8) to the mouse genome, and differential expression analysis was performed (total count filter cut-off = 5.0). The results were normalized to reads per million, and Gene-e (Broad Institute) was used to generate heat maps. The datasets generated in the current study are available from the corresponding author upon reasonable request.

### RNA extraction and real-time quantitative PCR analysis

Total RNA was extracted using Total RNA Kit II (Omega Bio-tek) according to the manufacturer’s instructions. cDNA was synthesized from 1 μg of total RNA using an RNA reverse transcriptase kit (Takara). Real-time qPCR was performed using the ABI PRISM 7000 Sequence Detection System (Applied Biosystems), and SYBR Premix Ex Taq II (Takara) was used for amplification according to the manufacturer’s instructions. The cDNA inputs were standardized, and PCR was performed for 40 cycles. The mouse organ tissues were homogenized to detect the virus gene copy number by qPCR (TaqMan Universal Master Mix II with UNG, 4440044, Applied Biosystems). The sequences of the primers are listed in Table 2.

**Table 2.**
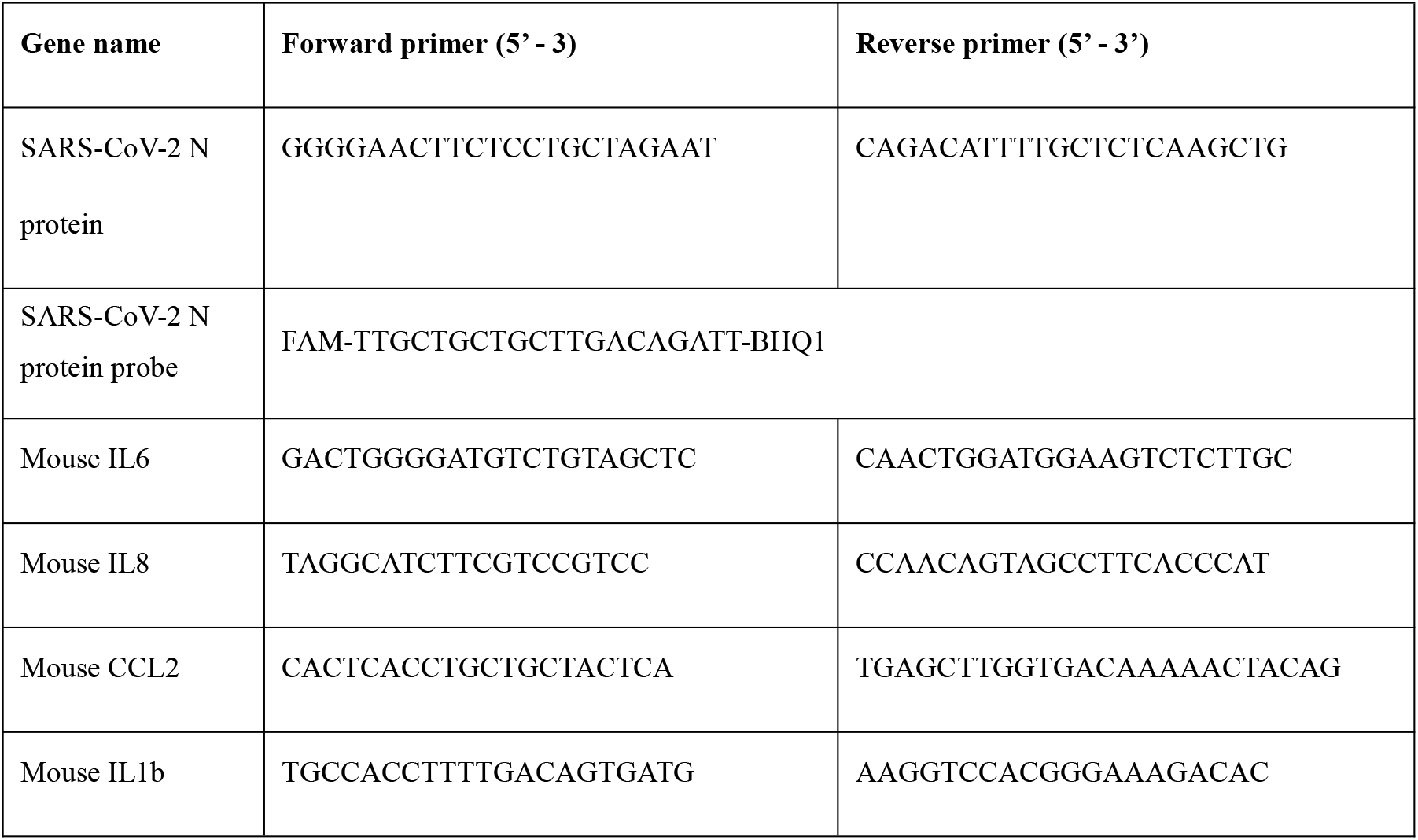
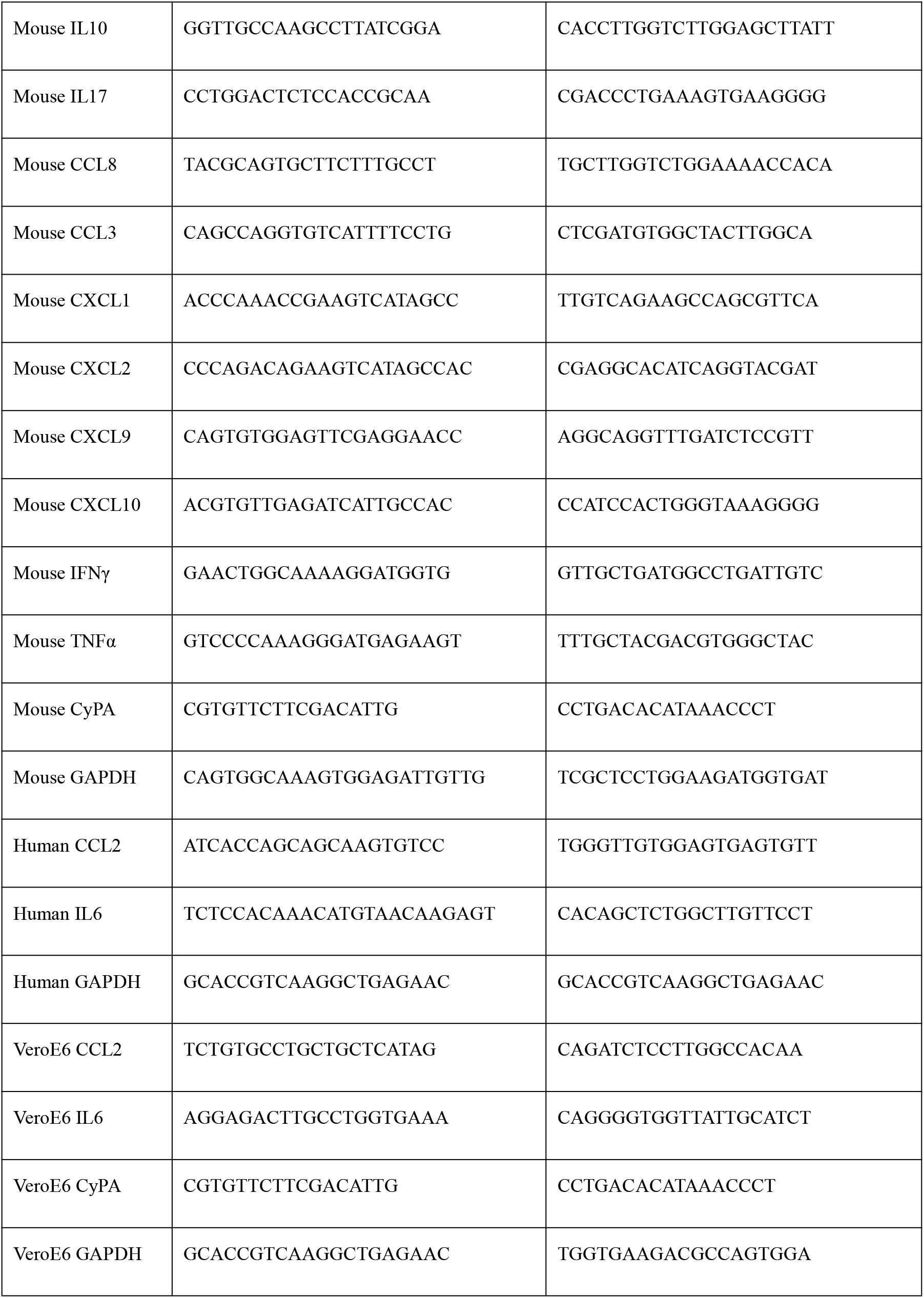
The primers for qPCR.

### Surface plasmon resonance (SPR)

SPR analysis was performed on a Biacore 3000 system (Biacore). In brief, His-CyPA (produced by our laboratory) was fixed to the surface of CM5 sensor chips (GE Healthcare Bio-Sciences AB) by an Amine Coupling Kit (GE Healthcare, BR-1000-50). The interaction between CyPA and CD147 was detected using the kinetic analysis/concentration series/direct binding mode; the flow rate was set to 15 μl/min, and the binding and dissociation time was 3 min. The same methods were applied to detect the interactions between CD147 and the SARS-CoV-2 spike proteins with mutations (N501Y, E484K, Y453F and N439K) (SinoBio, 40592-V08H8, 40592-V08H84, 40592-V08H80, 40592-V08H14) except that His-CD147 (produced by our laboratory) was fixed to the CM5 sensor chip surface. The results were analysed by BIAevaluation software (Biacore) to determine the affinity constants.

### Co-immunoprecipitation (Co-IP)

Co-IP assays were performed using a Pierce® Co-Immunoprecipitation Kit (26149, Thermo Fisher Scientific, Inc.) according to the manufacturer’s protocol. A mouse anti-human CyPA antibody (50 μg) and mouse anti-human CD147 antibody (Jiangsu Pacific Meinuoke Biopharmaceutical Co. Ltd, China, 50 μg) were used for antibody immobilization. The eluted proteins were detected by Western blot. Having boiled for 5 min, the eluted proteins were loaded onto a 12% SDS-PAGE gel and then transferred to a PVDF membrane (Millipore). After blocking with 5% nonfat milk for 1 h, the membrane was incubated with the corresponding primary antibodies at 4°C overnight. The images were developed following incubation with the secondary antibody (goat anti-mouse IgG (H+L), 31430, Thermo Fisher Scientific, 1:5000 dilution) at room temperature for 1 h.

### Transmission electron microscopy (TEM)

The lung tissues were cut into 1 mm^3^ pieces and fixed with PLP Fixing Solution (G2220, Solarbio) for at least 24 h. After washing with PBS, the sections were treated with osmic acid for 1.5 h. Then, the tissues were dehydrated with alcohol at gradient concentrations and soaked in acetone for 15 min. After embedding and polymerization overnight with epoxy resin, the slices were cut with an ultrathin slicer and glued to a perforated copper grid. Finally, each section was stained with lead citrate and uranium solution for 10 min, washed with water and dried. The ultrastructure was observed by transmission electron microscopy (JEM-1230, JEOLLTD).

### *In vitro* virus infection test

The cells were cultured at 37°C under 5% CO_2_ overnight, and the cell medium was then replaced by medium containing the virus. After the cells were infected for 1 h at 37°C, the virus supernatant was discarded, and the cells were washed twice with PBS. Finally, the cells were cultured with 2% FBS maintenance medium with or without CD147 antibody (Meplazumab, 15 or 60 μg/ml). At 48 h after infection, viral and cellular RNA were extracted together and detected by RT-qPCR.

### *In vitro* SARS-CoV-2 pseudovirus infection test

The SARS-CoV-2 pseudovirus and its variants (N501Y, N501Y-D614G, B.1.1.7 and B.1.351) expressing luciferase were obtained from the Institute for Biological Product Control, National Institutes for Food and Drug Control (Beijing, China). The SARS-CoV-2 pseudovirus was added to VeroE6 cells at a TCID_50_ of 1300 either with or without 60 μg/ml CD147 antibody. Meplazumab was used for inhibition assay of B.1.1.7, B.1.351, B.1.525, B.1.526(S477N), B.1.526(E484K), P.1 and P.2 variants, and Mehozumab and anti-CD147 antibody C were used for inhibition assay of B.1.617.1 and B.1.617.2 variants. Meplazumab, Mehozumab and anti-CD147 antibody C were humanized CD147 monoclonal antibodies, produced by Jiangsu Pacific Meinuoke Biopharmceutical Co. Ltd. Human IgG was used as a control. The luciferase signal was detected using a Dual-Luciferase Reporter Assay System (E1960, Promega) according to the manufacturer’s protocol.

### Statistical analysis

The cytokines detected by the protein chip were analysed by moderated *t*-statistics. ELISA data were measured using a parameter logistic curve. All multiplex fluorescence and immunohistochemical staining images were analysed by HALO Image Analysis Software, and significant differences were calculated by unpaired *t* tests. Correlations were analysed by Pearson correlation coefficients. Significant differences in cell and mouse tests were analysed by unpaired *t* tests with a two-tailed distribution. p < 0.05 was considered to be statistically significant. Statistical analyses were performed using GraphPad Prism, version 8.0.

---

**Figure S1.**
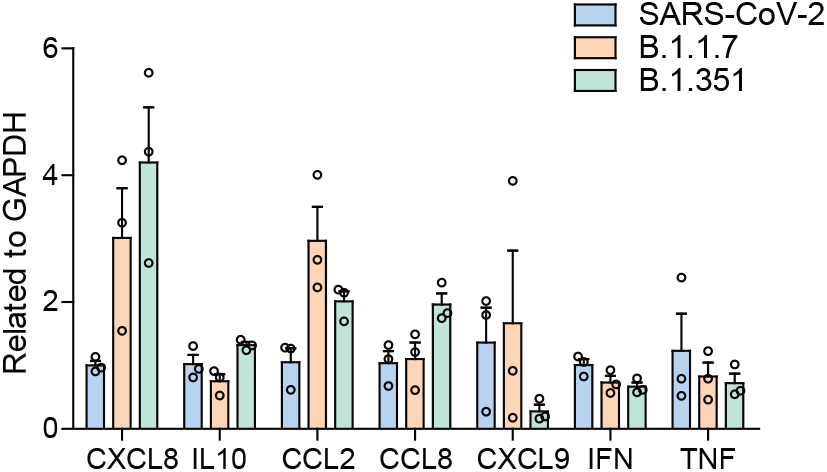
CD147 antibody can effectively inhibit the infection of SARS-CoV-2 and its variants. SARS-CoV-2, B.1.1.7 and B.1.351 virus infected VeroE6 cells for 48 h, and RNA was collected.The gene expression of the cytokines and chemokines determined by RT-qPCR. Gapdh is used as a reference gene (n = 5).

**Figure S2.**
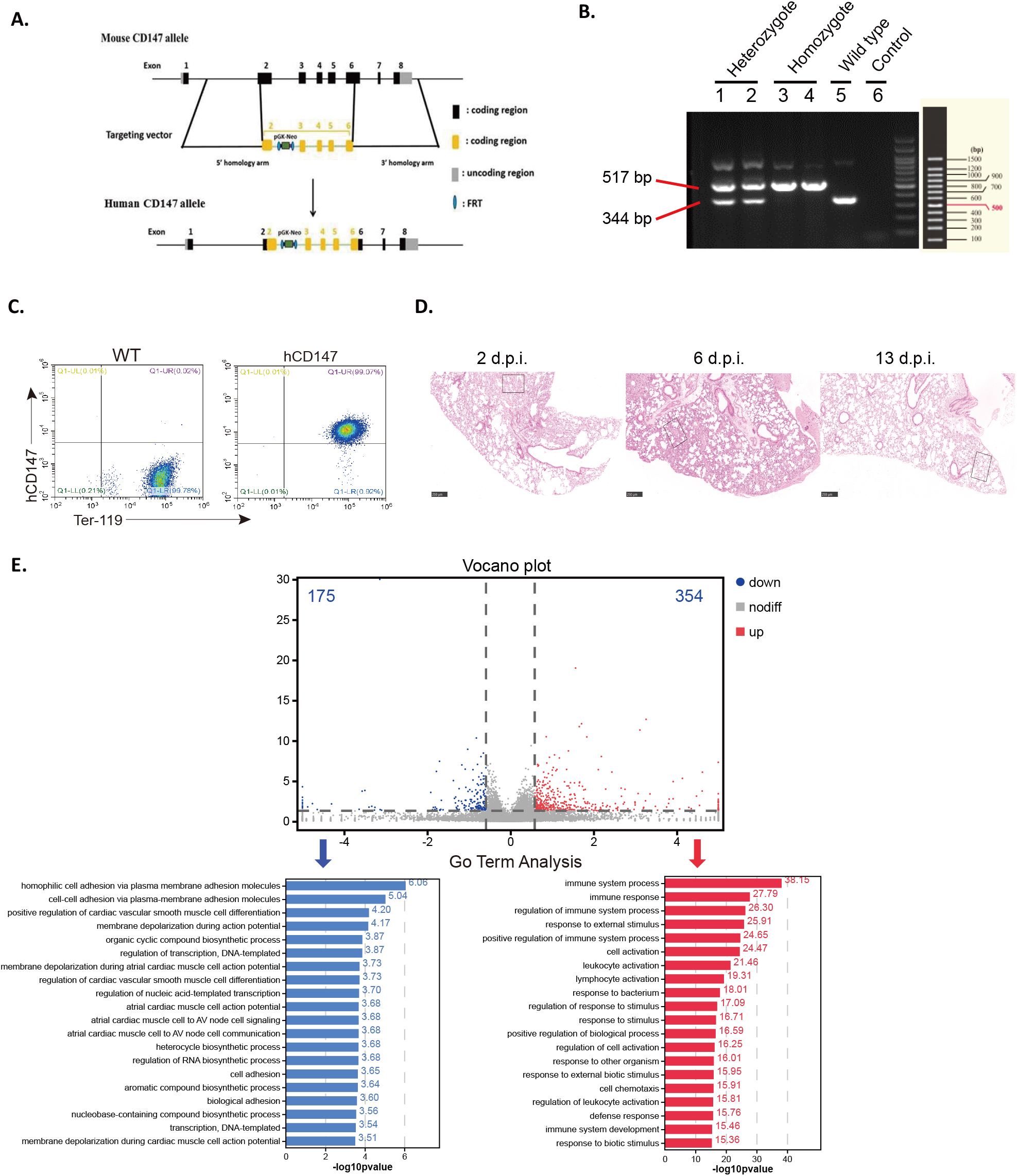
A hCD147 mouse model with susceptibility to SARS-CoV-2 infection and an inflammatory phenotype is established. **(A)** Schematic diagram of the method to construct hCD147 mice. (8) Identification of wildtype C57BL/6J and heterozygote and homozygote of hCD147 mice by nucleic acid electrophoresis. **(C)** Detection of human CD147 expression in peripheral blood of hCD147 mice by flow cytometry. H&E staining of lung sections from hCD147 mice after intranasal infection at 2, 6 and 13 d.p.i. (scale bars, 250 μm). **(E)** Volcano plots comparing differentially expressed genes in lung homogenates of C57BL/6J and hCD147 mice at 2 d.p.i. Red and blue indicate upregulated and downregulated genes, respectively, with a fold change > 1.5 and a false discovery rate (FDR) < 0.05. Gene Ontology (GO) enrichment analysis of biological process (BP) terms enriched in the upregulation and downregulation genes respectively.

**Figure S3.**
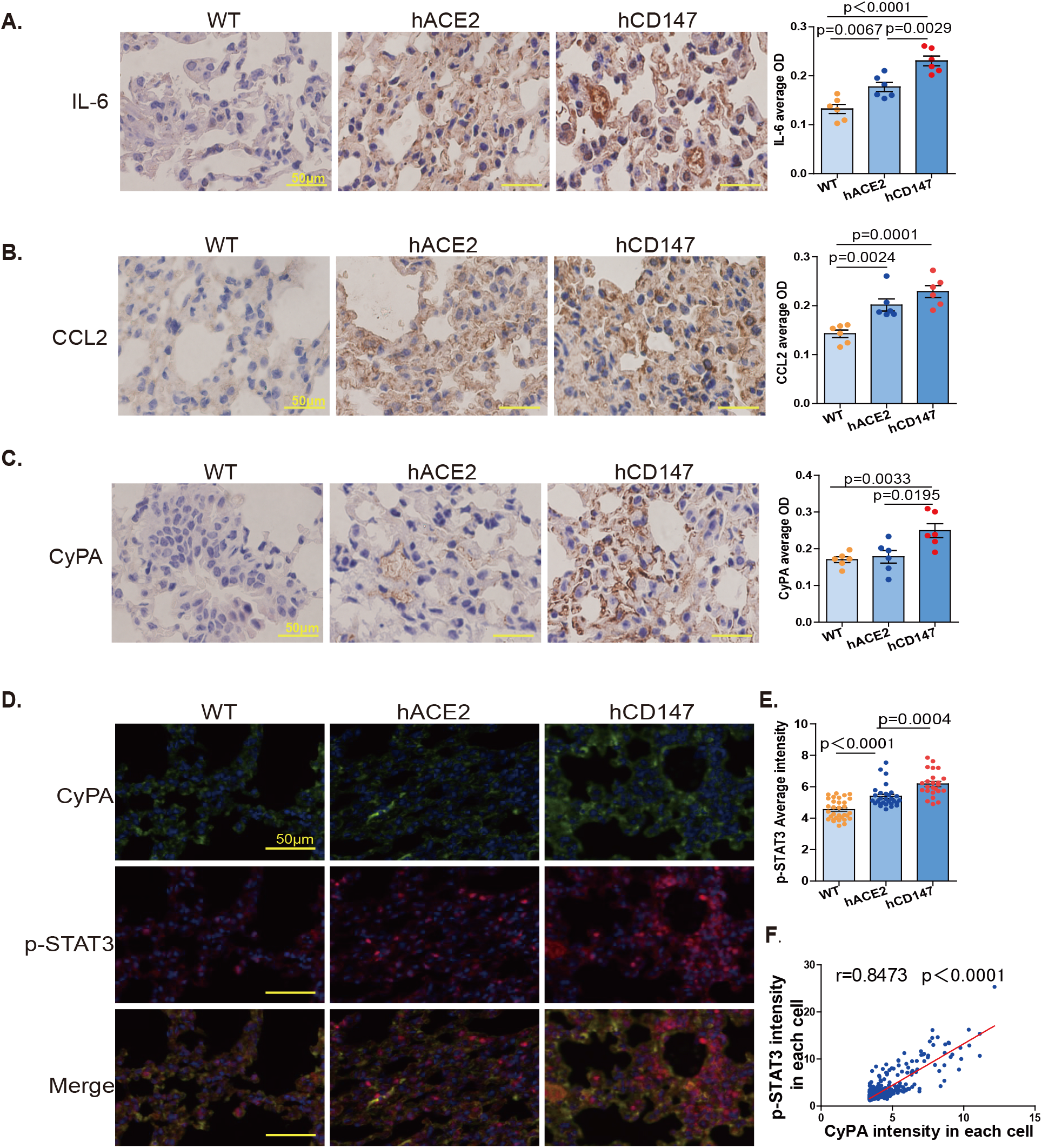
SARS-CoV-2 causes different pneumonia phenotypes and immune responses in the hACE2 and hCD147 mouse models. C57BU6J, hACE2 and hCD147 mice were intranasally inoculated with SARS-CoV-2 at a dose of 3 × 10^5^ TCID_50_. Lung tissues were collected at 2 d.p.i. for pathological analysis. **(A)** Detection of IL-6 expression by immunohistochemical staining. **(B)** Detection of CCL2 expression by immunohistochemical staining. **(C)** Detection of CyPA expression by immunohistochemical staining. **(D)** Detection of CyPA and p-STAT3 expression by multiplex immunofluorescence staining. **(E)** Statistics of average intensity of p-STAT3 in lung tissues of infected mice. **(F)** Correlation analysis between the intensity of p-STAT3 and the CyPA expression in the CyPA-positive cells in hCD147 mice.

**Figure S4.**
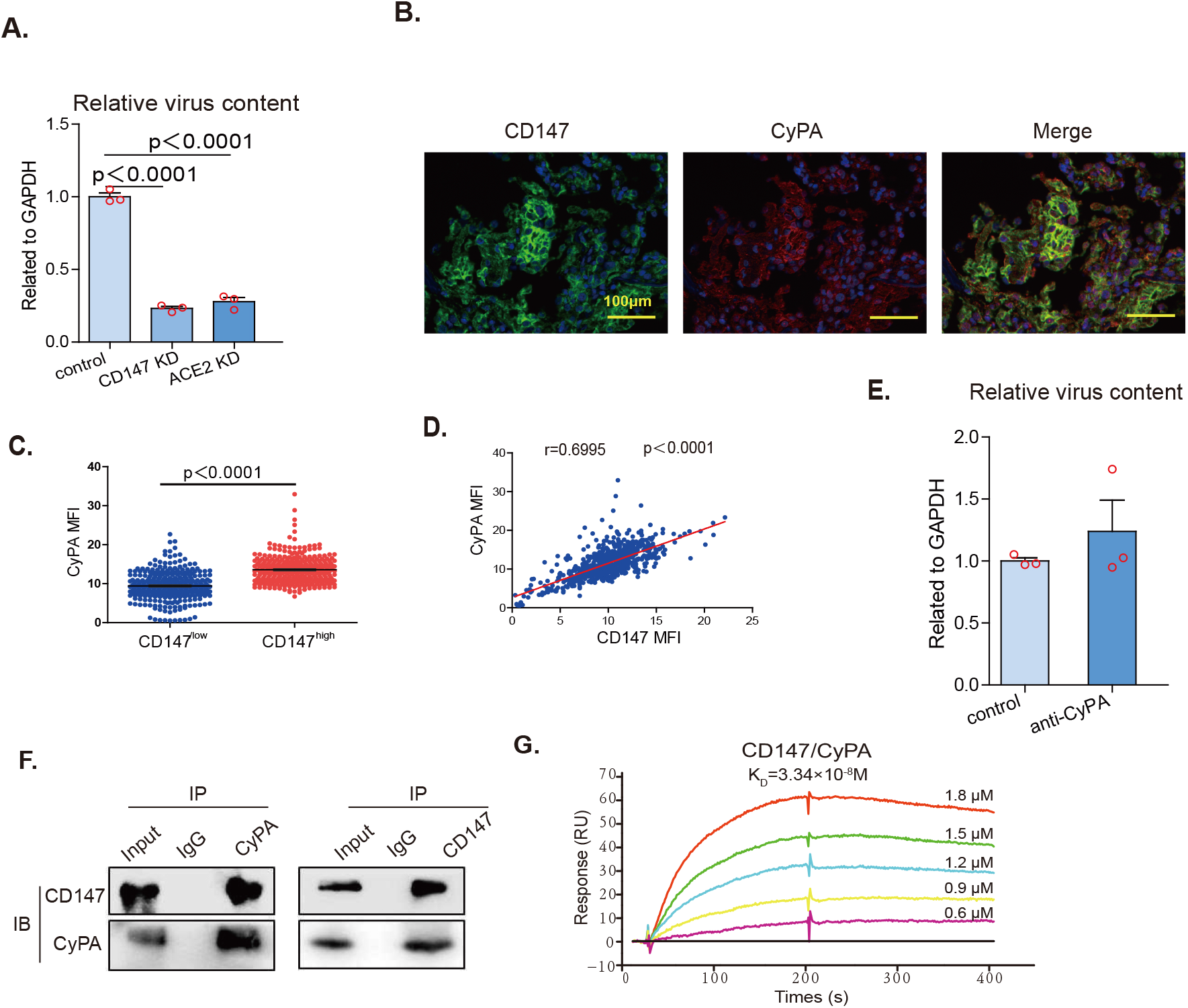
CyPA is a key intermediate proinflammatory factor involved in CD147-related immune responses. **(A)** CD147 or ACE2 in VeroE6 cells were knocked down by lentivirus, and then infected with SARS-CoV-2 for 48 h. The virus RNA was detected by RT-qPCR, normalized to Gapdh (n = 3, two-tailed unpaired t test). (8) Multiplex immunofluorescence staining was carried out for CyPA and CD147 (scale bars, 100μm). **(C)** Statistical difference of CyPA expression in CD147_low_ and CD147_high_ regions (Two-tailed unpaired t test). **(D)** Correlation analysis of CD147 intensity CyPA intensity. All the correlation scatter plots represent 13 cases of COVID-19 lung tissues data, which were analyzed by HALO software. All correlation analysis performed by Pearson correlation. The virus RNA was detected by RT-qPCR in VeroE6 cells infected with SARS-CoV-2 for 48 h, with or without CyPA antibody, normalized to Gapdh (n = 3, two-tailed unpaired t test). **(F)** The interaction between CyPA and CD147 was detected by Co-IP assay. **(G)** SP R analysis of the interaction between CyPA and CD147.

**Figure S5.**
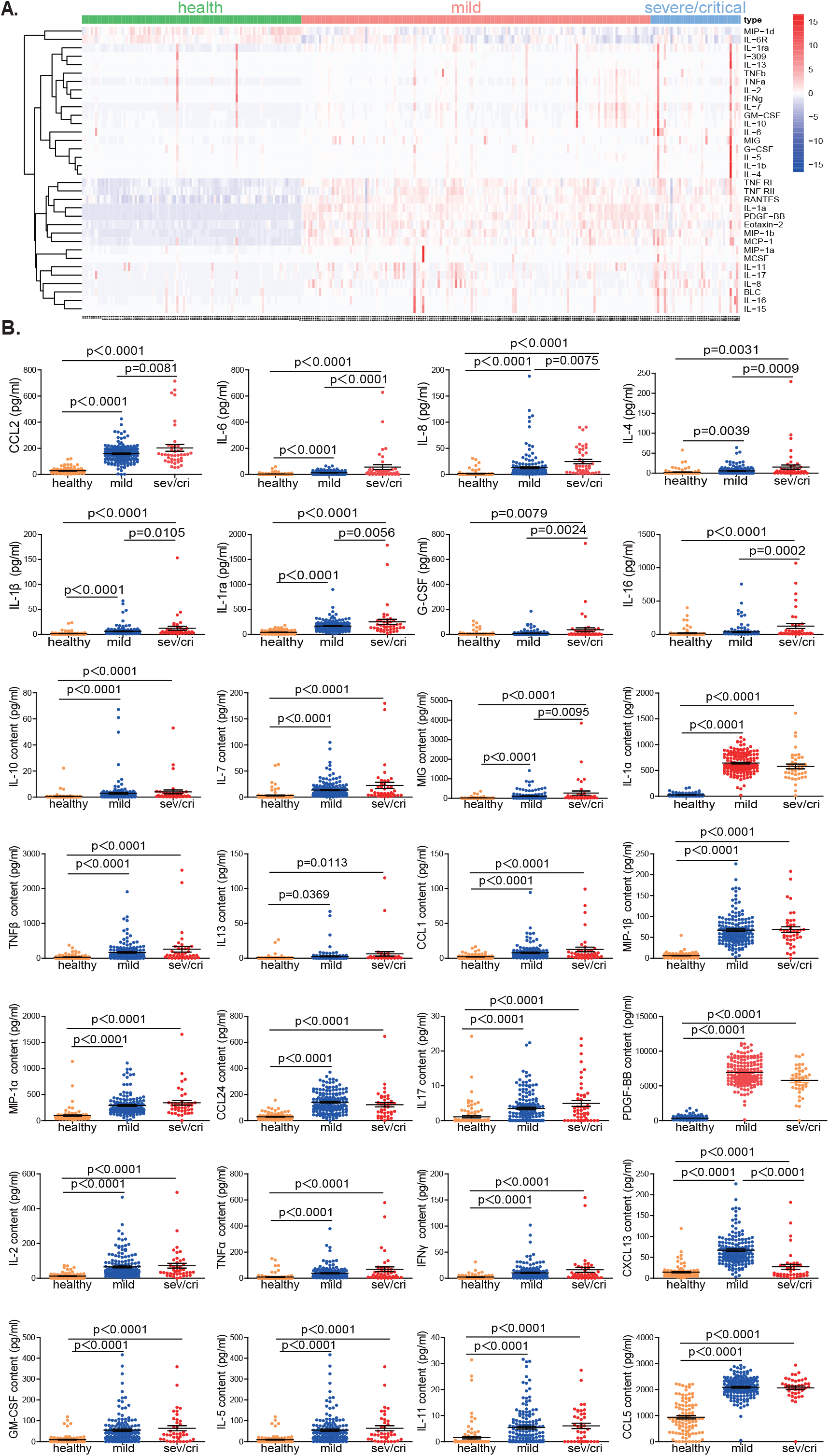
Human cytokines are detected in plasma from healthy individuals and COVID-19 patients. **(A)** Matrix analysis of 40 cytokines from healthy donors (n = 100) and COVID-19 patients with either mild (n = 159) or severe/critical (n = 41) disease. **(B)** Quantitative comparison of cytokines between healthy donors and COVID-19 patients with either mild or severe/critical disease (two-tailed unpaired *t* test).

**Figure S6.**
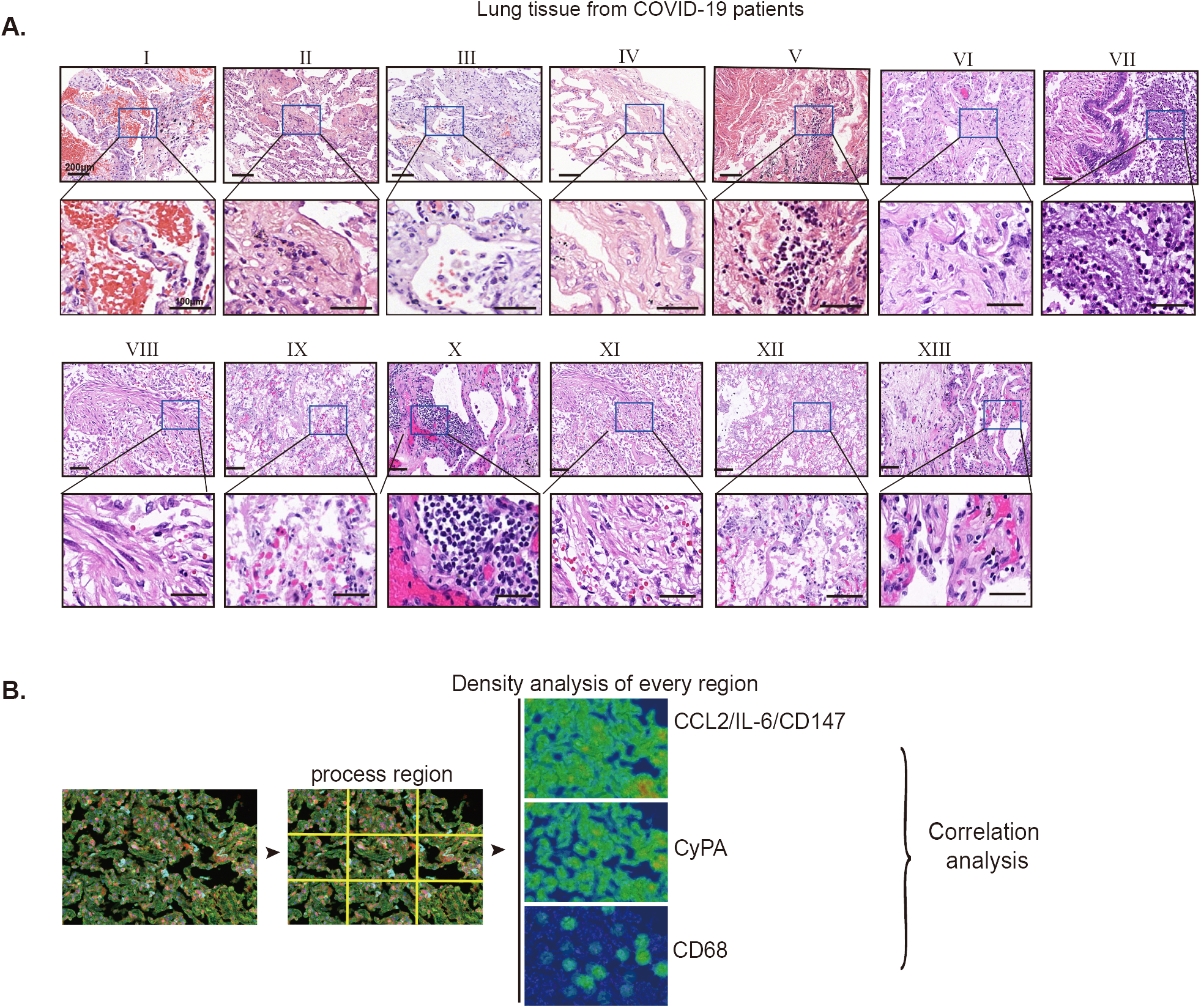
CyPA expression is positively correlated with the expression of CCL2 and IL-6 and infiltration of macrophages in the lung tissue of COVID-19 patients. **(A)** Pathological analysis of lung tissues from COVID-19 patients (n = 13). **(B)** The analytic process ofmultiplex immunofluorescence images. Each collected image was divided into process regions, and HALO image Analysis Software was used to analyze the MFI of each marker in each region.

**Figure S7.**
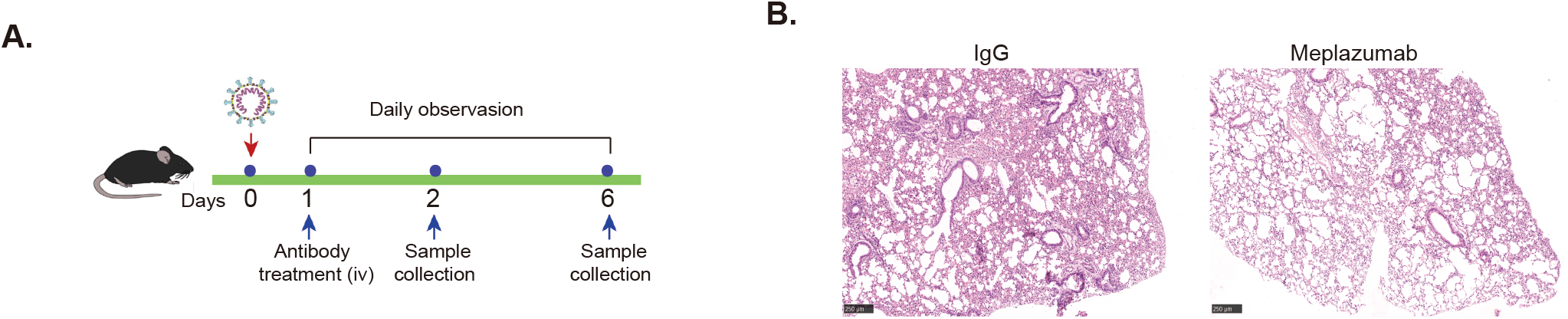
CD147 antibody reverses pulmonary inflammation caused by SARS-CoV-2. **(A)** Incubation of hCD147 mice via the intranasal at 3 × 10^5^ TCIO_50_ of SARS-CoV-2, and treatment with CD147 antibody the next day. Lung tissues were collected at 2 and 6 d.p.i. **(B)** H&E staining of lung sections from lgG and CD147 antibody group at 6 d.p.i. (scale bars, 250 μm).

## REFERENCES

1. P. Pymm et al., Nanobody cocktails potently neutralize SARS-CoV-2 D614G N501Y variant and protect mice. Proceedings of the National Academy of Sciences of the United States of America 118, (2021).

2. S. Jangra et al., The E484K mutation in the SARS-CoV-2 spike protein reduces but does not abolish neutralizing activity of human convalescent and post-vaccination sera. medRxiv, (2021).

3. M. Verghese et al., Identification of a SARS-CoV-2 Variant with L452R and E484Q Neutralization Resistance Mutations. J Clin Microbiol, (2021).

4. R. E. Chen et al., Resistance of SARS-CoV-2 variants to neutralization by monoclonal and serum-derived polyclonal antibodies. Nature medicine 27, 717–726 (2021).

5. J. Tian et al., Clinical characteristics and risk factors associated with COVID-19 disease severity in patients with cancer in Wuhan, China: a multicentre, retrospective, cohort study. Lancet Oncol 21, 893–903 (2020).

6. W. J. Guan et al., Clinical Characteristics of Coronavirus Disease 2019 in China. The New England journal of medicine 382, 1708–1720 (2020).

7. E. Barisione et al., Fibrotic progression and radiologic correlation in matched lung samples from COVID-19 post-mortems. Virchows Arch, (2020).

8. T. Mauad et al., Tracking the time course of pathological patterns of lung injury in severe COVID-19. Respir Res 22, 32 (2021).

9. M. A. Montero-Fernandez, R. Pardo-Garcia, Histopathology features of the lung in COVID-19 patients. Diagn Histopathol (Oxf), (2020).

10. J. Hadjadj et al., Impaired type I interferon activity and inflammatory responses in severe COVID-19 patients. Science 369, 718–724 (2020).

11. M. Tan et al., Immunopathological characteristics of coronavirus disease 2019 cases in Guangzhou, China. Immunology 160, 261–268 (2020).

12. F. Wang et al., Characteristics of Peripheral Lymphocyte Subset Alteration in COVID-19 Pneumonia. The Journal of infectious diseases 221, 1762–1769 (2020).

13. A. O. Hassan et al., A SARS-CoV-2 Infection Model in Mice Demonstrates Protection by Neutralizing Antibodies. Cell 182, 744–753 e744 (2020).

14. L. Bao et al., The pathogenicity of SARS-CoV-2 in hACE2 transgenic mice. Nature 583, 830–833 (2020).

15. N. Behloul, S. Baha, R. Shi, J. Meng, Role of the GTNGTKR motif in the N-terminal receptor-binding domain of the SARS-CoV-2 spike protein. Virus Res 286, 198058 (2020).

16. F. S. Oladunni et al., Lethality of SARS-CoV-2 infection in K18 human angiotensin-converting enzyme 2 transgenic mice. Nature communications 11, 6122 (2020).

17. R. Rathnasinghe et al., Comparison of transgenic and adenovirus hACE2 mouse models for SARS-CoV-2 infection. Emerg Microbes Infect 9, 2433–2445 (2020).

18. K. Wang et al., CD147-spike protein is a novel route for SARS-CoV-2 infection to host cells. Signal Transduct Target Ther 5, 283 (2020).

19. P. S. Arunachalam et al., Systems biological assessment of immunity to mild versus severe COVID-19 infection in humans. Science 369, 1210–1220 (2020).

20. T. S. Bonny et al., Cytokine and Chemokine Levels in Coronavirus Disease 2019 Convalescent Plasma. Open Forum Infect Dis 8, ofaa574 (2021).

21. Arunachalam, P.S., et al. Systems biological assessment of immunity to mild versus severe COVID-19 infection in humans. Science 369, 1210–1220 (2020).

22. S. H. Sun et al., A Mouse Model of SARS-CoV-2 Infection and Pathogenesis. Cell Host Microbe 28, 124–133 e124 (2020).

23. Dawar, F.U., et al., Potential role of cyclophilin A in regulating cytokine secretion. Journal of leukocyte biology 102, 989–992 (2017c).

24. X. Zhang, Y. Zhu, Y. Zhou, B. Fei, Interleukin 37 (IL-37) Reduces High Glucose-Induced Inflammation, Oxidative Stress, and Apoptosis of Podocytes by Inhibiting the STAT3-Cyclophilin A (CypA) Signaling Pathway. Med Sci Monit 26, e922979 (2020).

25. Glasmacher, E., et al. A genomic regulatory element that directs assembly and function of immune-specific AP-1-IRF complexes. Science 338, 975–980 (2012).

26. Johnson, G.L., and Lapadat, R. Mitogen-activated protein kinase pathways mediated by ERK, JNK, and p38 protein kinases. Science 298, 1911–1912 (2002).

27. Satoh, K., Nigro, P., and Berk, B.C. Oxidative stress and vascular smooth muscle cell growth: a mechanistic linkage by cyclophilin A. Antioxid Redox Signal 12, 675–682 (2010).

28. H. Bian et al., Meplazumab treats COVID-19 pneumonia: an open-labelled, concurrent controlled add-on clinical trial. medRxiv, (2020).

29. C. Y. Lee et al., Human neutralising antibodies elicited by SARS-CoV-2 non-D614G variants offer cross-protection against the SARS-CoV-2 D614G variant. Clinical & translational immunology 10, e1241 (2021).

30. F. Ali, A. Kasry, M. Amin, The new SARS-CoV-2 strain shows a stronger binding affinity to ACE2 due to N501Y mutant. Med Drug Discov 10, 100086 (2021).

31. S. Jangra et al., SARS-CoV-2 spike E484K mutation reduces antibody neutralisation. Lancet Microbe, (2021).

32. C. K. Yinda et al., K18-hACE2 mice develop respiratory disease resembling severe COVID-19. PLoS pathogens 17, e1009195 (2021).

33. D. Wu, X. O. Yang, TH17 responses in cytokine storm of COVID-19: An emerging target of JAK2 inhibitor Fedratinib. J Microbiol Immunol Infect 53, 368–370 (2020).

34. V. Vatsalya et al., Repurposing Treatment of Wernicke-Korsakoff Syndrome for Th-17 Cell Immune Storm Syndrome and Neurological Symptoms in COVID-19: Thiamine Efficacy and Safety, In-Vitro Evidence and Pharmacokinetic Profile. Front Pharmacol 11, 598128 (2020).

35. T. Shibabaw, Inflammatory Cytokine: IL-17A Signaling Pathway in Patients Present with COVID-19 and Current Treatment Strategy. J Inflamm Res 13, 673–680 (2020).

36. F. U. Dawar et al., Potential role of cyclophilin A in regulating cytokine secretion. Journal of leukocyte biology 102, 989–992 (2017).

